# Best Practices in Data Analysis and Sharing in Neuroimaging using MRI

**DOI:** 10.1101/054262

**Authors:** Thomas E. Nichols, Samir Das, Simon B. Eickhoff, Alan C. Evans, Tristan Glatard, Michael Hanke, Nikolaus Kriegeskorte, Michael P. Milham, Russell A. Poldrack, Jean-Baptiste Poline, Erika Proal, Bertrand Thirion, David C. Van Essen, Tonya White, B. T. Thomas Yeo

**Affiliations:** University of Warwick; McGill University; Heinrich-Heine University Düsseldorf; Otto-von-Guericke-University Magdeburg; MRC Cognition and Brain Sciences Unit; Child Mind Institute; Stanford University; University of California, Berkeley; Instituto Nacional de Psiquiatría Ramón de la Fuente Muñiz & Neuroingenia; Inria, Paris-Saclay University; Washington University in St. Louis; Erasmus University Medical Center; National University of Singapore

## Abstract

Neuroimaging enables rich noninvasive measurements of human brain activity, but translating such data into neuroscientific insights and clinical applications requires complex analyses and collaboration among a diverse array of researchers. The open science movement is reshaping scientific culture and addressing the challenges of transparency and reproducibility of research. To advance open science in neuroimaging the Organization for Human Brain Mapping created the Committee on Best Practice in Data Analysis and Sharing (COBIDAS), charged with creating a report that collects best practice recommendations from experts and the entire brain imaging community. The purpose of this work is to elaborate the principles of open and reproducible research for neuroimaging using Magnetic Resonance Imaging (MRI), and then distill these principles to specific research practices. Many elements of a study are so varied that practice cannot be prescribed, but for these areas we detail the information that must be reported to fully understand and potentially replicate a study. For other elements of a study, like statistical modelling where specific poor practices can be identified, and the emerging areas of data sharing and reproducibility, we detail both good practice and reporting standards. For each of seven areas of a study we provide tabular listing of over 100 items to help plan, execute, report and share research in the most transparent fashion. Whether for individual scientists, or for editors and reviewers, we hope these guidelines serve as a benchmark, to raise the standards of practice and reporting in neuroimaging using MRI.

## 1. Introduction

In many areas of science and in the public sphere there are growing concerns about the reproducibility of published research. From early claims by John loannidis in 2005 that “most published research findings are false” [Ioannidis2005] to the recent work by the Open Science Collaboration, which attempted to replicate 100 psychology studies and succeeded in only 39 cases [OpenScienceCollaboration2015], there is mounting evidence that scientific results are less reliable than widely assumed. As a result, calls to improve the transparency and reproducibility of scientific research have risen in frequency and fervor.

In response to these concerns, the Organization for Human Brain Mapping (OHBM) released “OHBM Council Statement on Neuroimaging Research and Data Integrity” [^1^ http://www.humanbrainmapping.org/OHBMDataIntegrity2014.] in June 2014, at the same time creating the Committee on Best Practices in Data Analysis and Sharing (COBIDAS). The committee was charged with (i) identifying best practices of data analysis and data sharing in the brain mapping community, (ii) preparing a white paper organizing and describing these practices, and (iii) seeking input from the OHBM community before (iv) publishing these recommendations.

COBIDAS focuses on data analysis and statistical inference procedures because they play an essential role in the reliability of scientific results. Brain imaging data is complicated because of the many processing steps and a massive number of measured variables. There are many different specialised analyses investigators can choose from, and analyses often involve cycles of exploration and selective analysis that can bias effect estimates and invalidate inference [Kriegeskorte2009, Carp2012].

Beyond data analysis, COBIDAS also addresses best practices in data sharing. The sharing of data can enable reuse, saving costs of data acquisition and making the best use of scarce research funding [^2^ The Lancet’s series on “Research: increasing value, reducing waste”,http://www.thelancet.com/series/research.] [Macleod2014], In addition, data sharing enables other researchers to reproduce results using the same or different analyses, which may reveal errors or bring new insights overlooked initially (see, e.g., [LeNoury2015]). There is also evidence that data sharing is associated with better statistical reporting practices and stronger empirical evidence [Wicherts2011], In short, data sharing fosters a scientific culture of transparency.

While many recent publications prescribe greater transparency and sharing of data (see, e.g., a pair of editorials in *Science* & *Nature* [Journals2014,McNutt2014]), such works are general to all of science or do not focus on human neuroimaging specifically (though see [Poline2012,Poldrack2014]). Thus the purpose of this paper is to elaborate some principles of open and reproducible research for the areas of practice relevant to the OHBM community. To make these principles practical and usable, we created explicit lists of items to be shared (Appendix D).

Working closely with OHBM Council, a “Proposed” version of this document was prepared by COBIDAS and first posted for comment in October 2015. Comments were collected via email and the COBIDAS blog [^3^ See http://www.humanbrainmapping.org/cobidas.] and a revised document [^4^ The present document differs from that voted on only in formating changes, typographical corrections, and the update of this particular sentence.] presented to the membership for an up/down vote in May 2016; 96% positive ballots were received. We note that while best practice white papers like this are not uncommon (see, e.g., [Alsop2014,Kanal2013,Gilmore2013]), they are generally authored by and represent the consensus of a small committee or at most a special-interest section of a larger professional body. Hence we are excited to present this work with the explicit participation and support of the OHBM membership.

### 1.1. Approach

There are different responses to the perceived crisis of reproducibility, with some simply letting the problem `self-correct’ as reviewers and readers become more aware of the problem, to more transformative measures like using blinded analyses or preregistration. In a blinded analysis all preprocessing, modelling and results generation is conducted with experimental labels hidden or scrambled, only being revealed after the analysis is fixed [MacCoun2015], while in preregistration all research hypotheses and analysis plans are published before data are collected, [Nosek2014].

The pragmatic approach behind this report is to increase the transparency of how research has been executed. Such transparency can be accomplished by comprehensive sharing of data, research methods and finalized results. This both enables other investigators to reproduce findings with the same data, better interrogate the methodology used and, ultimately, makes best use of research funding by allowing reuse of data.

The reader may be daunted by the sheer scale and detail of recommendations and checklists in this work (Appendix DV However we expect that any experienced neuroimaging researcher who has read a paper in depth, and been frustrated by the inevitable ambiguity or lack of detail, will appreciate the value of each entry. We do not intend for these lists to become absolute, inflexible requirements for publication. However they are the product of extensive deliberation by this panel of experts, and represent what we considered most effective and correct; hence, deviations from these practices may warrant explanation. Finally, we hope these lists can serve as tools for reviewers, e.g. as a reference for the importance of these items, and for editors and publishers, who can consider how this necessary detail can be conveyed without stringent word limits.

### 1.2. Scope

While the OHBM community is diverse, including users of a variety of brain imaging modalities, for this effort we focus exclusively on MRI. This encompasses a broad range of work, including task-based and resting-state functional MRI (fMRI), analyzed voxel-wise and on the surface, structural and diffusion MRI, but excludes other widely used methods like PET, EEG & MEG. We found that practice in neuroimaging with MR can be broken into seven areas that roughly span the entire enterprise of a study: (1) experimental design reporting, (2) image acquisition reporting, (3) preprocessing reporting, (4) statistical modeling, (5) results reporting, (6) data sharing, and (7) reproducibility.

Reproducibility has different and conflicting definitions (see Appendix B), but in this work we make the distinction between reproducing results *with the same data* [Peng2011] versus replicating a result *with different data* and possibly methods. Hence while this entire work is about maximizing replicability, the last section focuses specifically on reproducibility at the analysis-level.

This paper is structured around these areas, and for each we explore both general principles of open and reproducible research, as well as specific recommendations in a variety of settings. As the respective titles imply, for experimental design, data acquisition and preprocessing, studies are so varied that we provide general recommendations without recommending particular practices. Thus these sections focus mostly on thorough reporting and little on best practice. In contrast, for statistical modeling there are areas like task fMRI where mature methodology allows the clear identification of best practices. Likewise for the areas of data sharing, replication and reproducibility we focus exactly on those emerging practices that need to become prevalent.

We ask that authors challenge themselves: “If I gave my paper to a colleague, would the text and supplementary materials be sufficient to allow them to prepare the same stimuli, acquire data with same properties, preprocess in a similar manner and produce the same models and types of inferences as in my study?” This is an immense challenge! The purpose of this work is to guide researchers towards this goal and to provide a framework to assess how well a study meets this challenge.

## 2. Experimental Design Reporting

### 2.1. Scope

In this section we consider all aspects of the planned and actual experimental manipulation of the subject. This includes the type and temporal ordering of stimuli, feedback to be recorded and any subject-adaptive aspects of the experiment. It also encompasses basic information on the experiment such as duration, number of subjects used and selection criterion for the subjects. It is impossible to prescribe the “right” design for all experiments, and so instead the focus is on the complete reporting of all facets of the design.

### 2.2. General Principles

For experimental design, the goal of reproducible research requires the reporting of how the subjects were identified, selected, and manipulated. This enables a critical reader to evaluate whether the findings will generalize to other populations, and facilitates the efforts of others to reproduce and replicate the work.

### 2.3. Lexicon of Fmri Design

While other areas of these guidelines, like MRI physics and statistical modeling, have rather well defined terminology, we find there is substantial variation in the use of experimental design terms used in fMRI publications. Thus Box 2.1 provides terminology that captures typical use in the discipline. Since the analysis approach is dependent on the fMRI design, providing accurate and consistent characterization of the design will provide greater clarity.

There is often confusion between block and mixed block/event designs [Petersen2012], or block designs composed of discrete events. Thus we recommend reserving the term “block design” for paradigms comprised of continuous stimuli (e.g. flashing checkerboard) or unchanging stimuli presented for the entire length of a block (generally at least 8 seconds). All other designs comprise variants of event-related designs and must have their timing carefully described.

##### Box 2.1. fMRI Terminology

**Session.** The experimental session encompasses the time that the subject enters the scanner until they leave the scanner. This will usually include multiple scanning runs with different pulse sequences, including structural, diffusion imaging, functional MRI, spectroscopy, etc.

**Run.** A run is a period of temporally continuous data acquisition using a single pulse sequence.

**Volume.** A volume (or alternatively “frame”) is single 3-dimensional image acquired as part of a run.

**Condition.** A condition is a set of task features that are created to engage a particular mental state.

**Trial.** A trial (or alternatively “event”) is a temporally isolated period during which a particular condition is presented, or a specific behavior is observed.

**Event.** The term “trial” and “event” are often interchangeable. However, in certain situations of ‘compound-trials,’ a trial will consist of multiple subunits; for example, a working memory task may contain subunits of encoding, delay, and retrieval. In these cases the subunits are labeled as “events” and the “trial” is defined as the overarching task.

**Block.** A block (or alternatively “epoch”) is a temporally contiguous period when a subject is presented with a particular condition.

### 2.4. Design Optimization

Especially with an event-related design with multiple conditions, it can be advantageous to optimize the timing and order of the events with respect to statistical power, possibly subject to counterbalancing and other constraints [Dale1999, Wager2003], It is essential to specify whether the target of optimization is detection power (i.e. ability to identify differences between conditions) or estimation efficiency (i.e. ability to estimate the shape of the hemodynamic response, which requires jittering) [Liu2001]. It is likewise advisable to optimize your designs to minimize the correlation between key variables. For example, in model-based or computational fMRI experiments, variables such as reward, prediction error and choices will usually be highly correlated unless the design has been tuned to minimise this dependence. Be sure to include all possible covariates in a single statistical model to ensure variance is appropriately partitioned between these variables.

### 2.5. Power Analysis

The positive predictive value—the probability that an alternative hypothesis is true given a significant test—depends on the power of the study [Ioannidis2005], and underpowered studies have been found to been endemic in neuroscience as a whole [Button2013], Power analysis for imaging is difficult as the outcome is typically the entire brain image, and not a single univariate measure [Mumford2012]. However there are power analysis tools available to account for intra- and inter-subject fMRI variance at each voxel [Mumford2008] [^5^ http://fmripower.org.], as well as tools that account for the spatial structure of the signal [Joyce2012, Durnez2014] [^6^ https://sourceforge.net/projects/powermap & https://neuropower.shinyapps.io/neuropowerrespectively.]. Researchers hence can and should provide a realistic power computation that corresponds to the primary analysis of your study.

### 2.6. Subjects

The population from which the subjects are sampled is critical to *any* experiment, not just those with clinical samples. Be sure to note any specific sampling strategies that limited inclusion to a particular group (e.g. laboratory members, undergraduates at your university).

One should take special care with defining a “Normal” vs. “Healthy” sample. Screening for lifetime neurological or psychiatric illness (e.g. as opposed to “current”) could have unintended consequences. For example, in older subjects this could exclude up to 50% of the population [Kessler2005] and this restriction could induce a bias towards a ‘super healthy,’ thus limiting the generalization to the population.

### 2.7. Behavioral Performance

The successful execution of a task is essential for interpreting the cognitive effects of a task. Report behavioral measures in and out of the scanner, measures that are appropriate for the task at hand (e.g. response times, accuracy). For example, provide statistical summaries over subjects like mean, range and/or standard deviation. Note any pre-scan training, the setting (e.g. mock scanner vs. bench) and any training goals.

## 3. Acquisition Reporting

### 3.1. Scope

This section concerns everything relating to the manner in which the image data is collected on each subject. Again we do not attempt to prescribe best MRI sequences to use, but focus on the reporting of acquisition choices.

### 3.2. General Principles

Research can only be regarded as transparent when the reader of a research report can easily find and understand the details of the data acquisition. This is necessary in order to fully interpret results and grasp potential limitations. For the work to be reproducible, there must be sufficient detail conveyed to actually plan a new study, where data collected will have, e.g., similar resolution, contrast, and noise properties as the original data.

More so than many sections in this document, MRI acquisition information can be easily organized in ‘checklist’ form (see Appendix D). Thus in the remainder of this section we only briefly review the categories of information that should be conveyed.

### 3.3. Device Information

One fundamental aspect of data is the device used to acquire it. Thus every study using MRI should report basic information on the scanner, like make and model, field strength, and details of the coil used, etc.

### 3.4. Acquisition-Specific Information

Each acquisition is described by a variety of parameters that determine the pulse sequence, the field of view, resolution, etc. For example, image type (gradient echo, spin echo, with EPI or spiral trajectories; TE, TR, flip angle, acquisition time), parallel imaging parameters, use of field maps, and acquisition orientation are all critical information. Further details are needed for functional acquisitions (e.g. volumes per run, discarded dummy volumes) and diffusion acquisitions (e.g. number of directions and averages and magnitude and number of b-values [Jones2012]).

### 3.5. Format for sharing

While there is some overlap with Section 7. Data Sharing, there are sufficient manufacturer- and even model-specific details that we consider here related to data format. When providing acquisition information in a manuscript keep in mind that readers may use a different make of scanner, and thus you should minimize the use of vendor-specific terminology. To provide comprehensive acquisition detail we recommend exporting vendor-specific protocol definitions or “exam cards” and provide them as supplementary material.

When primary image data are being shared, a file format should be chosen that provides detailed information on the respective acquisition parameters (e.g. DICOM). If it is impractical to share the primary image data in such a form, retain as much information about the original data as possible (e.g. via NIfTI header extensions, or “sidecar” files). However, sensitive personal information in the acquisition metadata should be carefully removed through appropriate anonymization procedures before sharing (see Section 6. Data Sharing).

## 4. Preprocessing Reporting

### 4.1. Scope

This section concerns the extensive adjustments and “denoising” steps neuroimaging data require before useful information can be extracted. In fMRI, the two most prominent of these preprocessing steps are head-motion correction and intersubject registration (i.e., spatial normalisation), but there are many others. In diffusion imaging, motion correction, eddy current correction, skull stripping, and fitting of tensors (least squares, ROBUST, etc.) or more complex diffusion models are also key steps.

### 4.2. General Principles

As with other areas of practice, good reporting here requires authors to clearly detail each manipulation done to the data before a statistical or predictive model is fit. This is essential for reproducibility, as the exact outcome of preprocessing is dependent on the exact steps, their order and the particular software used. Moreover, the vast array of reprocessing options gives ample opportunity for p-hacking [Simmons2011] and thus it is vital to constrain choices and establish fixed preprocessing protocols whenever possible.

### 4.3. Software Issues

*Software versions.* Different tools implementing the same methodological pipeline, or different versions of the same tool, may produce different results [Gronenschild2012], Thus ensure that the exact name, version, and URL of all the tools involved in the analysis are accurately reported. It is essential to provide not just the major version number (e.g., SPM12, or FSL 5.0) but indicate the exact version (e.g. SPM12 revision 6225, or FSL 5.0.8). The version of software interpreters, e.g. Matlab or Python, should also be included as well as compilation conditions when known. To avoid ambiguities on the tool name, consider adding a Research Resource Identifier (RRID) [^7^ http://www.force11.org/group/resource-identification-initiative & https://scicrunch.org/resources.] [Bandrowski2015] citation for each tool used in addition to reporting the version. RRIDs index everything from software to mouse strains, and provide a consistent and searchable reference.

*In-house pipelines & software.* When using a combination of software tools, be sure to detail the different functions utilized from each tool (e.g., SPM’s realign tool followed by FreeSurfer’s boundary-based registration; see Reproducibility section for more on pipelines). In-house software should be described in detail, giving explicit details (or reference to peer-reviewed citation with such details) for any processing steps/operations carried out. Public release of in-house software through an open code repository is strongly recommended (e.g. Bitbucket or Github).

*Quality control.* Quality control criteria, such as visual inspection and automated checks (e.g., motion parameters), should be specified. If automated checks are considered, metric and criteria thresholds should be provided. If data has been excluded, i.e., due to scrubbing or other denoising of fMRI time series or removal of slices or volumes in diffusion imaging data, this should be reported.

*Ordering of steps.* The ordering of preprocessing steps (e.g., slice time correction before motion correction) should be explicitly stated.

*Handling of exceptional data.* Sometimes individual subjects will have problems, e.g. with brain extraction or intersubject registration. Any subjects that require unique preprocessing operations or settings should be justified and explained clearly, including the number of subjects in each group for case-control studies.

## 5. Statistical Modeling & Inference

### 5.1. Scope

This section covers the general process of extracting results from data, distilling down vast datasets to meaningful, interpretable summaries. Usually this consists of model fitting followed by statistical inference or prediction. Models relate the observable data to unobservable parameters, while inference quantifies the uncertainty in the estimated parameter values, including hypothesis tests of whether an observed effect is distinguishable from chance variation. Inference can also be seen as part of making predictions about unseen data, from the same or different subjects. Note that while we make a clean distinction between preprocessing and modeling, there is some overlap (e.g. movement can be a preprocessing “correction” or part of a model) and they can in fact interact [Strother2002].

### 5.2. General Principles

For statistical modeling and inference, the guiding principle of openness dictates that the reader of published work can readily understand what statistical model was used to draw the conclusions of the paper. Whether accidental or intentional (i.e. for brevity), omission of methodological details needed to reproduce the analyses violates these principles. For maximal clarity, be sure to describe all data manipulation and modeling in the methods section [Gopen1990], For maximal transparency, report all regions of interest (ROIs) and/or experimental conditions examined as part of the research, so that the reader can gauge the degree of any HARKing (Hypothesizing After the Results are Known) [Kerri1998]; to absolutely minimize HARKing, register your hypothesis and analysis before exploring the data.

### 5.3. Assumptions

Every modelling and inference method described below makes assumptions about the data analyzed. Take the time to understand assumptions and the implications on your results. Just to name a few for linear models: The correct model (no missing variables), additive model (no missing interactions), normality of errors (no outliers), etc. [Luo2003], In group analyses, pay special attention to reversal paradoxes where an effect can be flipped if an important covariate is omitted [Tu2008], And always take care when removing outliers, as informative observations may be discarded. Consider using the analysis that has the weakest assumptions while still providing the needed inference, e.g. tools like permutation or bootstrap (e.g. [Eklund2015,Bellec2010].

### 5.4. Software

See the previous section for details on how to describe the exact software and pipeline used.

### 5.5. Mass Univariate Modelling

A simple univariate model fit to each voxel or surface element is known as a mass univariate modelling approach, and is an essential tool for everything from task fMRI, structural MRI measures like Voxel Based Morphometry, scalar diffusion measures like Fractional Anisotropy or even resting state fMRI, when measured with low frequency variance (see *Other Resting-State Analyses* below). Regardless of the type of data, a mass-univariate linear model is specified by five types of information: Dependent variables, independent variables, model, estimation method and inference method (where inference refers to quantification of uncertainty of estimated parameters and hypothesis testing).

While the *dependent variable* (or response) may be unambiguous (e.g. for T2* BOLD), be sure to identify it in any nonstandard analysis. Itemize each *independent* (or explanatory) *variable* in each model used. In a first level fMRI model, this includes the usual condition effects, as well as motion regressors added to explain nuisance variation. In a second level or group model, independent variables include the group assignment (e.g. patient vs. control) or other between-subject effects that may or may not be of interest (e.g. age or sex). Report non-trivial contrasts, linear combinations of independent variables, that are used to interrogate the experimental effect of interest. Variables generally do not need to be centered [^8^ http://mumford.fmripower.org/mean_centering.], but do indicate how and if this was done.

While software may make the *model* and *estimation method* seem ‘automatic’, a short description is needed for a complete scientific report. See Appendix C for examples of short descriptions of commonly used task fMRI models. Beyond the mass univariate model, there is growing use of other types of models, including local multivariate, whole-brain multivariate, etc. Regardless of the model, be sure to note the essential details of the estimation procedure.

The *inference method* is used to flag some voxels or elements as “active” or “different”, as distinguished from background noise, and is a crucial final step. In brain imaging, inference usually amounts to a thresholding procedure, though if ROIs are used, it should also include computation of confidence or credible intervals. ROIs can be defined based on anatomy or previous literature [Poldrack2007], or based on functional results with great care and attention to circularity (see *Extracted Data* below).

While the dangers of multiple testing are generally well understood, one review found only 59% of studies used a correction for multiple testing [Carp2012], While there are domains where stringent corrections are not suitable (e.g. presurgical fMRI, where false positive risk has to be balanced against the dangers of missing eloquent tissue), every study needs to address the issue, using a standard correction method or defending the lack of correction. Thus clearly state the type of inference and the manner of multiple-testing correction or reasons for no correction. Note that the inference method description “5% cluster wise inference” doesn’t specify the cluster-forming threshold nor the multiple-testing correction measure (e.g. familywise or false discovery rate). Also describe the volume, sub-volume, or surface domain for which multiple-testing correction has been performed, and whether multiple volumes/ROIs have been corrected for.

### 5.6. Connectivity Analyses

Functional and effective connectivity encompass a broad range of methods, from data-driven multivariate or clustering methods on high resolution voxel-wise data, to highly structured physiological-based models on a small number of regions. Methods are still evolving for resting-state fMRI in particular, but careful execution of a study requires considering topics similar to task fMRI modeling: response variables, model, estimation method and inference method.

The goal of most connectivity analyses is to understand the relationships among multiple *response (dependent) variables.* These variables can be defined by regions-of-interest (ROIs), in which case be sure to report the number of ROIs and how the ROIs are defined (e.g. citable anatomical atlas; auxiliary fMRI experiments). State whether analyses were carried out as a voxelwise whole-brain analysis or by using cortical surfaces or CIFTI ‘grayordinates’ (surface vertices + subcortical gray matter voxels [Glasser2013]). For seed-based analyses, or small-scale (e.g. Bayes Net) methods, provide the rationale and method for selecting the particular ROIs. Carefully describe how time series were attributed to each ROI (e.g. averaging, median, or eigenvariate), and detail any additional (temporal or spatial) filtering or transformations (e.g. into wavelet coefficients) used, or nuisance variables (e.g. motion parameters) ‘pre-regressed’ out of the data.

A number of exploratory multivariate methods are used to understand high-dimensional fMRI data in a lower dimensional space. These include Principal Component Analysis, Multidimensional Scaling, Self Organizing Maps, and Independent Component Analysis (ICA), of which ICA is probably the most widely used. For any such method report the model variant (e.g. spatial or temporal ICA), the estimation method (i.e. algorithm) and the number of dimensions or components used and, crucially, how this number was selected. ICA fitting and interpretation depends on choices about scaling (a.k.a. variance normalization), both to the data before fitting and as a constraint between spatial and temporal components; describe the type of scaling (i.e. variance normalization) applied to data and extracted components. Always report how the components were sorted and selected for analysis, whether for the primary analysis or as part of a denoising procedure; if using a post-hoc task regression model supply associated task model details (see above).

As any nuisance variation jointly influencing multiple voxels/regions can be mistaken for brain connectivity, it is essential that careful preprocessing has been applied, including artifact removal (See Section 3).

For many connectivity analyses the *model* is nothing more than the summary measure of dependence, e.g. Pearson’s (full) correlation, partial correlation, mutual information, etc. However, be sure to note any further transformations (e.g. Fisher’s Z-transform, regularization of partial correlation estimates). For seed-based analyses, describe the voxel-wise statistic or regression model (and other covariates) used. For regression-based group ICA analyses (“dual regression”, or “PCA-based back-reconstruction”), clearly describe how the per-subject images are created. As with task-fMRI, any group analysis should be described in terms of dependent variables, independent variables, model, estimation method, and inference method. For graph analysis methods based on binary connection matrices, state how thresholding was done and consider the sensitivity of your results to the particular threshold used. While lag-based methods like Granger have been criticised for fMRI [Smith2012], they remain suitable for EEG and MEG.

For functional connectivity, *inference* typically focuses on making statements comparing two or more groups of subjects or assessing the impact of a covariate. Ensure that it is clear what is the response being fed into the group model. For some connectivity analyses, like Structural Equation Modelling or Dynamic Causal Modelling, the inference concerns selecting among a set of models. Be sure to justify and enumerate the models considered and how they were compared; describe how evidence for model selection was aggregated over multiple subjects. Discuss the prior distributions used and their impact on the result. For graph-based analyses, detail the construction of adjacency matrices (i.e. what was binarized and how), or if using weighted measures, how the weights are computed. Note the problems of comparing networks of different size or overall connection density [vanWijk2010].

### 5.7. Other Resting-State Analyses

The analysis of resting state data does not necessarily incorporate connectivity. Methods like Amplitude of Low Frequency Fluctuations (ALFF) [Zang2007] and fractional ALFF (fALFF) [Zou2008] summarise brain activity with absolute (ALFF) or relative (fALFF) BOLD variance, and Regional Homogeneity (ReHo) [Zang2004] measures local consistency of signals. These methods produce a map per subject that can be analyzed with a mass univariate model (see above).

### 5.8. Multivariate Modelling St Predictive Analysis

Predictive methods focus on estimating an outcome for each experimental trial, block or subject, often using multivariate models. Multivariate methods exploit dependencies between many variables to overcome the limitations of mass univariate models, often providing better explanatory or predictive models. In brain imaging, predictive methods are often called decoding or multi-voxel pattern analyses [Norman2006]; an example of a multivariate analysis is representational similarity analysis [Kriegeskorte2008]. A complete description should include details of the following: Target values, features, predictive model, and training and validation method.

The *target values* are the outcomes or values to be predicted, which may be discrete or continuous. It should be made clear exactly what is being predicted, and what are the relative frequencies of this variable (e.g. proportions in each group, or a histogram for a continuous target). Unbalanced group sizes are not a problem but require appropriate evaluation measures, as described in the next section.

The *features* are the variables used to create the prediction, and often are not the raw data themselves but derived quantities. In addition, some features may be discarded in the process of feature selection. If feature selection is based on the target values there will be a tendency to over-fit, and then feature selection must be embedded in the validation framework (see below). It is essential that the analysis pipeline is described in sufficient detail to capture the definition of each element of the feature, any feature selection that precedes model-training, and any feature transformations (including possible standardization).

The *predictive model* is the type of method used to map features to targets. Typical examples include linear discriminant analysis and support vector machines. The model is distinct from the algorithm or training procedure used to optimize the parameters of the method (i.e. usually to minimize prediction error on held-out data). Be sure to clearly identify the model used and (if used) the specific machine learning library used.

Finally, the *training and validation method* is perhaps the most important facet of a predictive analysis. This comprises the algorithm used to build the predictive model and the framework used to evaluate the model. Training may be nothing more than fitting a regression model, but more typically consists of a complex algorithm that depends on the tuning of hyper-parameters. In the validation step the model’s predictive performance (e.g. accuracy) is assessed using an independent dataset or a cross-validation framework (e.g. leave-one-out, k-fold cross-validation, stratified cross validation). Clearly specify the algorithm used, what objective function was optimized, how the algorithm’s convergence was established (for iterative methods), and any post-processing of the fitted model. Be sure to clearly describe how hyper-parameters were estimated, including the choice of the hyper-parameter grid, figure-of-merit optimized, the type of validation scheme used, and the use of an averaging strategy to produce a final classifier. In particular, identify which hyper-tuning parameters were optimized outside vs. inside a cross-validation loop: The reported accuracy is evidently valid only if *all* estimated hyper-parameters are optimized inside the loop as part of a nested cross-validation procedure with three-way splits providing disjoint training, testing and validation datasets, and no information from the test data enters the optimization of any of the parameters.

## 6. Results Reporting

### 6.1. Scope

The reporting of statistical results is inextricably tied to the statistical modeling and inference procedures of the previous section. However, a scientific investigation invariably requires dozens of analyses, inferences and views of the data, and thus any published report typically contains a subset of all output of every statistical procedure completed. Thus we feel that results reporting deserves its own section here, providing guidance on how authors should select and present the outcomes of the modeling process.

### 6.2. General Principles

Transparency of published research requires that the reader can easily interpret the results shown and, crucially, what results were considered but then not shown. Unreported selective inference (a type of file drawer effect) inflates the significance of results shown and will stymie efforts to replicate a finding.

### 6.3. Mass Univariate Modelling

For reporting single univariate outcomes, like average BOLD response in an ROI or global mean FA, there is a wealth of best practice guidelines available [^9^ http://www.equator-network.org.] [Altman2008]. For mass univariate models, there are four general classes of information that need to be carefully described: Effects tested, tables of brain coordinates, thresholded maps, parcellated maps, and extracted data.

A complete itemization of the *effects tested* must be presented, identifying the subset that are presented. This is necessary to understand the true magnitude of the multiplicity involved and the potential danger of selection biases. For example, if a study has a multifaceted design allowing various main and interaction effects to be considered, effects tested and omitted should be enumerated, including references to previously published results on the current dataset. A full sense of how extensively the data has been explored is needed for the reader to understand the strength of the results.

*Tables of coordinates* historically have often been the *only* quantification of the results, and now should be complemented with sharing of full statistic images (see, e.g., NeuroVault [^10^ http://neurovault.org.]). If coordinates are reported, each table or sub-portion of a table should be clearly labeled as to what contrast / effect it refers to (nature of the contrast, individual versus group result, group size), and should have columns for: Anatomical region, X-Y-Z coordinate, T/Z/F statistic, and the P-value on which inference is based (e.g. voxel-wise FWE corrected P; or cluster-wise FDR corrected P); if cluster-wise inference is used, the cluster statistic (e.g. size, mass, etc) should be included. Avoid having multiple columns of results, e.g. multiple XYZ columns, one for increases, one for decreases, or one for left hemisphere, one for right hemisphere.

The table caption should clearly state (even if in repetition of the body text) the significance criterion used to obtain these coordinates, and whether they represent a subset of all such significant results (e.g. all findings from whole-brain significance, or just those in a selected anatomical region). If T or F statistics are listed, supply the degrees of freedom. Whenever possible, provide effects sizes at the selected coordinates together with 95% confidence intervals. Finally, the space (i.e., Talairach, MNI, fsaverage) of the coordinate system should be noted.

The *thresholded map figures* perhaps garner the most attention by readers and should be carefully described. In the figure caption clearly state the type of inference and the correction method (e.g. “5% FWE cluster size inference with P=0.001 cluster-forming threshold”), and the form of any sub-volume corrections applied. For small volume or surface ROI corrections, specify whether or not the ROI was identified prior to any data analysis and how it was defined. Always annotate threshold maps with a color bar for the statistic values; when showing multiple maps, use a common color bar when feasible; and always indicate right and left. Avoid common fallacies in interpreting maps; e.g. an activation in region A but not region B doesn’t mean A is significantly more active than B [Poldrack2008], and lack of activation is not evidence of no activation. Most important, publicly share the original statistic images, unthresholded and thresholded, so readers can explore the maps themselves in 3D (see Data Sharing below).

*Extracted data* from images aids the interpretation of the complex imaging results, and is presented as effect magnitudes, bar plots, scatter plots or activation time courses. Computed from a single voxel/vertex, or an average or principal component of a set of voxels/vertices, they however present a great risk for “circularity” [Vul2009; Kriegeskorte2009]. Specifically, when the voxels summarized are selected on the basis of a statistic map, they are biased estimates of the effect that map describes. Thus it is essential that every extracted summary clearly address the circularity problem; e.g. “derived from independently-formed ROI”, or “values based on voxels in a significant cluster and are susceptible to selection bias”. When working with single regions and uncorrected P-values, consider the current discussions on the limitations of P-values [Wasserstein2016] and in particular how P=0.05 can amount to very weak evidence of an effect [Nuzzo2014].

### 6.4. Functional Connectivity

The critical issues when reporting functional connectivity differ between types of approaches, for example exploratory multivariate vs. seed-based correlation methods, which provide whole maps, versus confirmatory multivariate methods for a handful of regions.

When reporting multivariate decomposition methods like ICA or PCA, state how the number of components were selected. Wth either ICA or seed-based analyses, when conducting inference on multiple networks, be sure to account for multiplicity when searching over the networks. For example, if testing for patient vs. control differences in the default mode, attentional, visual and motor networks, the inference must account for not only the voxels within networks, but additionally for searching the four IC maps for significance.

### 6.5. Multivariate Modelling St Predictive Analysis

While it may appear that predictive analyses are trivial to report (“Accuracy was X%, p(X> chance)<0.001”), there are in fact two broad types of information to convey: Evaluation & interpretation.

*Evaluation* refers to the assessment of a fitted classifier on out-of-sample data. As shown in the tabular listing, there are several measures of classifier performance that should be reported aside from overall accuracy (percentage of correct predictions). For example, when group sizes are unequal, be sure to also report average or balanced accuracy (accuracy per group, averaged).

Do not make claims of “above chance accuracy” unless based on confidence intervals or some appropriate formal test, ideally a permutation test [Combrisson2015]. For regression report prediction R^2^, though be aware this may be negative when the explained variance is low (but is not necessarily truly zero). Avoid using a correlation coefficient as an evaluation metric (computed between actual and held-out-predicted continuous values) as this is susceptible to bias [Hastie2011, Ch7].

*Interpretation* of the fitted classifier allows potential insights to brain function or structure that drives prediction, though must be done with care (see e.g. [Haufe2014]). In particular, be sure not to over-interpret whole brain weight maps as localizing the source of decoding information, as the very multivariate nature of the method means it is impossible to isolate a single region as being responsible for classification. Voxels or vertices containing significant information may receive small or zero weight if a regularisation penalty is used in fitting. Conversely, voxels/vertices with high absolute weight may contain little predictive signal, but may mostly serve to cancel correlated noise, thus improving classifier performance. The same caveats apply in the context of an encoding model that predicts brain responses from various experimental features; e.g. a predictor with large weight may be cancelling effects of other predictors and may not by itself contain any information about the voxel in question. Solutions to this problem include adding relevant (e.g. smoothness, continuity) priors to the multivariate model to improve its interpretability, and using resampling techniques like *stability selection* to enhance the reliability of the estimated classifier weights [Varoquaux2012], Mapping procedures that conduct the same analysis at every location, such as multivariate searchlight mapping, can also outline regions that are predictive in isolation of activity elsewhere and thus complement whole-brain classification methods.

Finally, just as mass univariate analyses can be weakened by ‘data dredging’ through scores of contrasts, a predictive analysis is also less meaningful if it is the (say) 10th analytical approach tried on a single dataset. It is essential to itemize the analyses attempted, both to convey what doesn’t work and the size of the model space considered.

## 7. Data Sharing

While previous sections have largely described good practice that is (more or less) prevalent in the community, this and the next section concerns practices that are currently scarce. Thus these sections are necessarily more prescriptive, providing explicit suggestions on ways to change how we conduct studies, meeting the challenges of making neuroimaging science as transparent and reproducible as possible.

### 7.1. Scope

Neuroimaging, relative to other disciplines like genetics and bioinformatics, has lagged behind in widespread acceptance of data sharing. This section outlines the practicalities of sharing of data and results, including issues related to the use of software infrastructure, data repositories and the details surrounding retrieval of data.

### 7.2. General Principles

Data sharing is one of the cornerstones of verifiable and efficient research, permitting others to reproduce the results of a study and maximizing the value of research funds already spent. However, to fully realize this value, data should not just be “available on request”, but shared in a data repository that is well organized, properly documented, easily searchable and sufficiently resourced as to have good prospects for longevity. In this respect, we support the FAIR data guiding principles according to whom data should be Findable, Accessible, Interoperable and Re-usable [^11^ https://www.force11.org/node/6062.]. To fulfill these goals, there are four key elements to a successful data sharing effort: planning, databases, documentation, & ethics.

### 7.3. Planning for Sharing

Data sharing is most onerous when done as an afterthought [Halchenko2015], Instead, if data sharing is considered when a study is planned and initiated as part of a complete data management plan, the additional effort required will be minimal. A data sharing plan should establish a viable path for any qualified researcher to gain access to the data. Planning efforts should begin with ethics (consent forms) and funding agencies, by creating realistic funding roadmaps for long-term stewardship of data. Having realistic workflows and a proper technical infrastructure are important prospective steps. In addition, there are a number of considerations and hurdles for publishers, each of which have their own policies for data sharing.

A key to an effective data sharing is the use of a strict naming structure for files and directories. This regularity brings a number of benefits, including greater ease in finding errors and anomalies. But most valuable, organized data facilitates extensive use of scripting and automation, reducing time needed for analysis and quality control (QC). Best practice is to use a standardized data structure; for example, the recently developed BIDS standard [^12^ http://bids.neuroimaging.io.] provides a detailed directory hierarchy for images and a system of plain text files for key information about a study’s data. This structure is used by OpenfMRI [^13^ http://openfmri.org.], making it easy to upload data to that repository. Whatever the system, arranging your data in a regular structure will simplify all efforts to manipulate and—specifically—share your data.

Another essential decision to make early in a study is exactly what kinds of data are to be shared. The exact data shared must be consistent with the ethics of the study (see below, Ethics). But once suitably anonymised, there are still the various versions of image data to choose from: DICOM files from the scanner for each subject; “raw” converted data (e.g. NIFTI), free of any preprocessing; ready-to-model fMRI data for each subject, having all of the basic processing completed; per-subject summary maps, e.g. one effect/contrast image per subject in fMRI; per-study statistic maps. Sharing raw data gives more options to other users (DICOM being the rawest), while sharing preprocessed images makes it easier for others to immediately start analyzing your data. Sharing of extensively processed data, such as (unthresholded) statistical maps and underlying structural data (e.g., volumes and cortical surfaces of individuals and/or group averages) can be very valuable, enabling readers of an article to access much more information than can be conveyed in a static image in a publication. Finally, share as many QC measures as possible as well as providing a PASS/FAIL summary for each dataset, allowing easy selection of usable data while allowing users to revisit QC decisions.

Decide at the outset with whom the data is to be shared and at what stage, as it may be useful to share data with collaborators prior to publication, then more freely after publication. We support the widest sharing of data possible, but in certain (e.g. clinical) circumstances this may not be possible without additional protective measures, such as an explicit Data Use Agreement (DUA), or the use of “data on request” services (e.g. Clinical Study Data Request [^14^ http://clinicalstudydatarequest.com.]) that enable assessment of compliance with such terms. Again consistent with ethics, have a data management plan that clearly specifies whether data can be freely distributed, or under exactly what constraints it can be shared. For example, in large-scale databases, data may be freely shared within a project, with some limits to other related projects, or with yet more constraints to the general public. Establishing these limits before a single subject is scanned will save many headaches down the road. Instead of setting the exact rules for data use yourself, consider using an established license, like from the Creative Commons [^15^ http://creativecommons.org/licenses.] or Open Data Commons [^16^ http://opendatacommons.org/licenses.], saving yourself time and making the terms of use clear to users.

For large-scale, multi-site studies, the greater effort put into harmonization of experimental paradigms, data acquisition, analysis and modeling, the easier it will be to amalgamate the data later. If separate databases are used, then an ontological standardization is important, establishing how to map data fields and the data dictionaries between sites.

Another facet to consider is the sharing of data analysis pipelines scripts and any provenance traces. These are generally free of ethical concerns (unless protected information like subject names creeps into a script!) and there is great value in allowing others to recreate your results and apply your methods to new data. This is discussed in greater detail below (see Documentation).

Finally, whenever possible use publication as the milestone for sharing. The longer you wait the harder it becomes to assemble all the pieces (data, scripts, etc), plus the article can then have the DOI/URL reference to the data.

In short, a comprehensive data management plan—that involves all authors, collaborators, funding agencies, and publishing entities—is essential no matter what is shared and should be considered from the outset of a study. Without such planning, in a jumble of folders and after a graduate student or post-doc has moved on, data can effectively be lost.

### 7.4. Databases

While a highly organized arrangement of data in a folder hierarchy is prerequisite for good data management, it does not in itself constitute a database. A database, in addition to organizing data, is searchable and provides access controls. Databases for imaging data may include non-imaging data and allow direct entry of data. There are a number of imaging-oriented databases, ranging in scale, complexity, features and, crucially, effort needed to install and maintain them. As individual users are unlikely (and not advised!) to create imaging databases, we review the considerations when choosing a database.

Consider access control options, and exactly who and which types of users should be allowed to enter data, and access the data. There may be some types of data (e.g. sensitive behavioral tests or essential personal information) that require special, restricted access. The ability to modify existing data should be highly restricted, ideally with a form of audit control that records the nature of the changes. A public repository must of course provide access for external users.

Comprehensive search functionality is important, especially for large scale, multi-project databases. Useful features include being able to select subsets of data of interest, e.g. finding subjects that have a certain age range, IQ and a clinical diagnosis, with two different imaging modalities. Once a selection is made, some systems may only let you download data, while others may provide quick visualization or extensive analysis options. Especially when working with large repositories, the availability of a scriptable query interface can be handy for complex queries.

Consider the ability of a system to handle heterogeneous data. Most imaging databases will accommodate the most basic demographic information, but may not accept more than one modality (e.g. both MRI and EEG) or other types of essential data, like clinical evaluations or batteries of psychological tests. Consider carefully all the data that comprises your studies and whether it can all be stored in one unified system. Some systems allow staff to directly enter subject information, and even conduct batteries of psychological tests on subjects, eliminating double entry and reducing the risk of errors.

Finally, assess the complexity of installation and maintenance of a system. At a single site, the system must be easy to install and maintain, while a database for a multi-site study will necessarily be more complex and require adequate expertise to manage. As part of this, ensure there is detailed documentation for maintainers, as well for end users on how to navigate the resource. And if serving as a repository, a database must additionally possess a long-term preservation plan.

Now, with a variety of mature imaging databases available, building a de novo home-grown database cannot be recommended. For example, IDA [Mueller2005], XNAT [Marcus2007], COINS [Scott2011], and LORIS [Das2012] are four established and well-resourced systems for longitudinal, multi-modal, web-based data storage and querying, with proper user control. Some of these tools interface to high performance computing platforms for mass processing (e.g. IDA to LONI [Dinov2009] or LORIS to CBRAIN [Das2016]) and can be an important element in reproducibility (see *Reproducibility* section).

While these established databases are becoming easier to install and maintain, we acknowledge that in low resource environments they may be impractical. In these settings, the use of highly structured storage of imaging data (see BIDS above) and extensive use of scripting is the best approach, and facilitates a transition later to a formal database. In most research environments, however, informatics support should be regarded as a necessity and funded accordingly, if for no other reason than to obtain the maximal value of the data collected, now and for years to come.

### 7.5. Documentation

Even an organized and searchable database is of no use, unless users have access to information describing what is actually stored in the repository. Clear documentation on the studies within a repository, the data acquisition and experimental paradigm detail are all examples of information that are needed to make use of information in a database. If processed data and results are stored, details on the preprocessing and models fit are also essential. The documentation should be written for a wide audience, including members from multiple disciplines. The extensive documentation for the Human Connectome Project [^17^ http://humanconnectome.org/documentation.] provides a great example of how to describe data (unprocessed and minimally preprocessed) as well as the acquisition and preprocessing methods in a large and complex database.

A form of self-documentation is provenance, i.e. recording exactly what happened to data through preprocessing and modeling. These “provenance traces” can help track-down problems and provide invaluable reference for others who want to replicate previous studies. While provenance is not usually recorded, the AFNI BRIK [^18^ http://afni.nimh.nih.gov/afni/doc/faq/39.] and MINC [^19^ http://www.bic.mni.mcgill.ca/software/minc.] formats have forms of provenance tracking, and the NIDM project [^20^ http://nidm.nidash.org.] is developing a framework to save this information in a standard format. Pipeline software like LONI Pipeline [^21^ http://loni.usc.edu/Software/Pipeline.] or Nipype [^22^ http://nipy.org/nipype.] explicitly provide such provenance records.

### 7.6. Ethics

There are ethical issues that promote and constrain data sharing. In favor of data sharing is the gain in knowledge from re-used data that can improve health in patient populations, not to mention how sharing can make better use of the public funds spent on research. But the data to be shared pertains humans whose rights and wishes must be respected. Hence suitably crafted ethics and consent documents are essential for data sharing. While in the United States de-identified data is currently not “protected health information” and should be able to be shared, regulations differ between countries and institutions and are subject to change. In the United Kingdom, a separate set of data protection laws exist and must be complied with. Thus be sure to consult your ethics or institutional review board, and any data protection office, before acquiring data with the intent of sharing, as well as before releasing data. The Open Brain Consent project [^23^ http://open-brain-consent.readthedocs.org.] can also be of use, providing sample forms written specifically to account for later sharing of data. Some level of anonymization will be required, ensuring all sensitive personal information is withheld or suitably coarsened or obscured (e.g. reporting only age in years at scan time instead of birth date), and/or applying a “de-facing” procedure to anatomical MRI images. Careful ‘scrubbing’ (e.g. removing subject names from DICOM files, or analysis pathnames) is required to ensure no personal information is discussed.

## 8. Reproducibility

We make the distinction articulated by [Peng2011] and others that *reproducible* results can be recreated by others using the same data and software as shared by the original authors, while a *replication* is the traditional scientific goal of independent researchers using independent data and possibly distinct methods to arrive at the same scientific conclusion (see Appendix B). While some have argued that reproducibility is secondary, and that “one should replicate the result not the experiment” [Drummond2009], recent failures to replicate high-impact results and occasional but acutely concerning examples of outright fraud have made the case for the importance of reproducibility.

### 8.1. Scope

We focus on analysis-level reproducibility, i.e. the ability to recreate the results of a well-defined analysis using the same data. All of the recommendations of this paper are in the service of the clear, unambiguous reporting of design, data and analysis workflow as recommended by the FAIR principles [^24^ https://www.force11.org/node/6062.] [Wilkinson2016], EQUATOR Network [^25^ http://www.equator-network.org.] [Altman2008], Reward Alliance [^26^ http://researchwaste.net.] and ARRIVE guidelines [^27^ https://www.nc3rs.org.uk/arrive-guidelines.] [Kilkenny2010], To further make your analysis as reproducible as possible, ensure it is *documented, archived* and *citable*.

### 8.2. Documentation

As even the operation system version can influence the exact results obtained [Glatard2015], be sure to cite the computational infrastructure as well as versions of software used.

When the analysis involves multiple tools, some formal description of the workflow connecting these tools should be provided. Tools such as BrainVISA [Cointepas2001], LONI pipeline [Rex2003], NiPype [Gorgolewski2011], PSOM [Bellec2012], Automatic Analysis Pipeline [Cusack2015], SPM batch [Penny2006] and CBRAIN [Das2015] may help structure and describe workflows. myExperiment [DeRoure2010] can be used to share and run workflows online (see for instance this FSL fMRI workflow from the LONI Pipeline environment [^28^ http://www.myexperiment.org/workflows/2048.]).

Any additional information on provenance will aid in efforts to reproduce your analysis. For example, tools like NiPype & the LONI Pipeline Processing environment [MacKenzie-Graham2008] records an exact “provenance trace” of the analysis, and the MINC [^29^ http://en.wikibooks.org/wiki/MINC/Reference/MINC2.0_Users_Guide.] and AFNI BRIK formats also store histories of analysis commands used to create a file. The Neuroimaging Data Model (NIDM [Keator2013]) is being actively developed to describe all steps of a data analysis in analysis-program-independent fashion.

Even when the data and workflow used in an analysis are properly documented, it may not be easy to reproduce the exact same data, for instance figures, as presented in a publication. Consider the use of literate programming tools such as iPython notebooks (used for instance in [Waskom2014]), or R-based Sweave [Leisch2002], Another example involves 'scene’ files that store all of the information (including links to the associated data files) that is needed to exactly reproduce a published figure. Currently, scene files are supported by the Connectome Workbench [Marcus2013] and Caret [VanEssen2001] software platforms.

### 8.3. Archiving

The analysis documentation should be archived in a long-term accessible location on the web. Of course, even with excellent documentation resources may disappear, become inaccessible, or change, further challenging reproducibility.

Open-source software is more likely to be available long term and is thus recommended. Whenever available, report on the availability of tools in repositories such as the INCF software center [^30^ http://software.incf.org.], the NITRC Resource Registry [^31^ http://www.nitrc.org.], or in software suites such as NeuroDebian [Halchenko2012] or Lin4Neuro [Nemoto2011].

A good way to facilitate reproducibility is to create and release a virtual machine (VM) or a container with the software and pipelines used in the analysis. A good starting point is the NeuroDebian VM [^32^ http://neuro.debian.net or https://hub.docker.com/_/neurodebian.] that can be further customized for a particular use case. Examples of other practical solutions that demonstrate this approach are the Nipype vagrant box and the NITRC Computational Environment [^33^ http://www.nitrc.org/plugins/mwiki/index.php/nitrc:UserGuide-NITRC_Computational_Environmen] (used e.g. in [Ziegler2014]), both NeuroDebian-based VMs, and Niak [^34^ http://simexp.github.io/niak.] (available on DockerHub) [^35^ http://hub.docker.com.]. Of course licensing may prevent creating comprehensive VM. With Matlab code, consider using the Matlab Compiler to create standalone applications or free alternatives such as GNU Octave.

### 8.4. Citation

URLs tend to “decay” over time [Habibzadeh2013], making them inappropriate to cite online material permanently. Instead, Digital Object Identifiers (DOIs) provide a persistent way to index digital data. Various platforms are now available to host your data and workflows and create DOIs for them, such as Zenodo [^36^ http://zenodo.org.], figshare [^37^ http://figshare.com.] or the Harvard Dataverse system [^38^ http://dataverse.harvard.edu.] (see examples in [Tustison2014], [Soelter2014] & [Holmes2015]). Data hosting projects may also register DOIs directly to organizations such as DataCite [^39^ http://www.datacite.org.] or CrossRef [^40^ http://www.crossref.org.].

## 9. Conclusions

In this work we have attempted to create an extensive (but not comprehensive) overview of reporting practices and, to a lesser extent, the practices themselves needed to maximize the openness and replicability of neuroimaging research. We have focused exclusively on MRI, but many of the suggestions and guidelines will easily translate to other areas of neuroimaging and related fields.

This document is inevitably dated by the current technology and means of reporting scientific results. As these evolve this document will need to be updated and revised. Updates and the current version of these guidelines will be available at the COBIDAS website [^41^ http://www.humanbrainmapping.org/cobidas.].

A reaction to the extensive checklists (Appendix D) could be “What human can put all that into their paper!?”, and our response is that ideally no human should, that is it should be a computer’s job. Many of the elements to report exist in some machine readable form, but in countless different forms. Thus the next important work to be done is to align these checklists to a controlled vocabulary, e.g. from NIH’s Common Data Elements [^42^ https://www.nlm.nih.gov/cde.]. Once terms are set like this, they are more easily entered into a format like ISA-Tab [^43^ http://www.isa-tools.org.], a table-based system for recording machine-readable metadata (now used, e.g., by the journal *Scientific Data),* and the stage is then set to develop the tools to automatically extract and export such vital meta-data.

## Acknowledgements

Please see Appendix A.

## Appendix A. COBIDAS Membership & Acknowledgements

Upon creating COBIDAS in June 2014, Dr. Nichols was named as chair and subsequently invited nominations from the OHBM membership. From over 100 nominees Dr. Nichols selected a dozen experts from the membership that reflected the diversity of OHBM, with the final list approved by Council. The different constituencies considered included: Researchers focusing in cognitive applications, clinical applications, methods and database developers; different geographic areas; sex; representation of junior researchers; and, to facilitate communication within OHBM leadership, at least one member from Council and one member from the OHBM Program Committee.

The full list of members is as follows (in alphabetical order).

Simon Eickhoff, Department of Clinical Neuroscience and Medical Psychology Heinrich-Heine University Dusseldorf, Dusseldorf, Germany.

Alan Evans, Montreal Neurological Institute, McGill University, Montreal, Canada.

Michael Hanke, Otto-von-Guericke-University Magdeburg, Germany.

Nikolaus Kriegeskorte, MRC Cognition and Brain Sciences Unit, Cambridge, UK Michael Milham, Child Mind Institute, New York City, USA.

Thomas Nichols (chair), University of Warwick, UK.

Russell Poldrack, Stanford University, Stanford, USA.

Jean-Baptiste Poline, University of California, Berkeley, Berkeley, USA.

Erika Proal, Instituto Nacional de Psiquiatria Ramon de la Fuente Muniz *&* Neuroingenia, Mexico City, Mexico.

Bertrand Thirion, Inria, Paris-Saclay University, France.

David Van Essen, Washington University, Division of Biology and Biomedical Sciences, St. Louis, USA.

Tonya White, Erasmus MC, Rotterdam, Netherlands.

BT Thomas Yeo, National University of Singapore, Singapore.

We are deeply grateful to guest members Tristan Glatard & Samir Das, informatics experts who contributed greatly to data sharing and reproducibility sections, and (then) OHBM President Karen Berman and former OHBM president Peter Bandettini, who participated in a number of calls.

We are indebted to Ben Inglis for allowing us to use his fMRI acquisition methods reporting checklist [Inglis2015] as a template for our Acquisition Reporting checklist. We are also grateful to the following for input on different specific sections: Ben Inglis, Doug Noll, Robert Welsh, Jorg Stadler, Derek Jones and Stamatios Sotiropoulos on the Data Acquisition section; Tim Behrens on the Experimental Design section; Vince Calhoun and Christian Beckmann on Statistical Modeling & Inference; Ged Ridgway for Preprocessing and Statistical Modeling & Inference; for help with Ting Xu for Statistical Modeling & Inference.

We would like to thank the members of the brain imaging community who took time to comment on the original draft of this work. Through email comments, marked up manuscripts and blog entries, we received over 100 comments, each of which has been considered and, with only a few exceptions, integrated into the present manuscript. In addition to anonymous participants in the blog, these contributors included:

David Abbott, Carsten Allefeld, John Ashburner, Mary Askren, Peter Bandettini, Andreas Bartsch, Patrick Bellgowan, Saskia Bollmann, Matthew Brown, Karen Davis, Simon Davis, Ekaterina Dobryakova, Simon Eskildsen, Satrajit Ghosh, Matt Glasser, Tristan Glatard, Enrico Glerean, Douglas Greve, Luis Hernandez-Garcia, Emilia lannilli, Tom Johnstone, Jorge Jovicich, Varsha Khodiyar, Vesa Kiviniemi, Thomas Liu, Martin Lotze, Dan Lurie, David Macfarlane, Clare Mackay, Paul Mazaika, Donald McLaren, Janaina Mourao Miranda, Jeanette Mumford, Helmut Nebl, Cyril Pernet, Aina Puce, Ged Ridgway, Alard Roebroeck, Tom Ross, Keith Schneider, David Soltysik, Stamatios Sotiropoulos, Stephen Strother, Paul Summers, Karsten Tabelow, Jussi Tohka, Robert Turner, Ilya Veer, Tor Wager, Alle Meije Wnk, Tal Yarkoni, and Yu-Feng Zang.

Thanks also to Alex Bowring for assistance in final preparation of this manuscript.

## Appendix B. Defining Reproducibility

A number of terms with overlapping meaning are used to refer to the merits of scientific findings, including reproducibility, replicability, and reliability. Here we attempt to set the terminology and clarify their meaning as used this report.

*Replication* is a cornerstone of the scientific method. A replication, where independent researchers use independent data and possibly distinct methods to arrive at the same original conclusion, is the ultimate standard for validating a scientific claim.

Roger Peng [Peng2011] suggested a specific notion of *reproducibility* in the computational sciences. He articulated a kind of reproducibility where independent researchers use the exact same data and code to arrive at the original result. Within this there is a spectrum of reproducibility practice, ranging from a publication sharing only code, or code and data, to the best case, where detailed scripts and code and data are shared that produces the results reported in the paper when executed.

The US Food & Drug Administration also has definitions to describe the precision of measurements, as codified by terms from the International Standards Organization (ISO), “repeatability” and “reproducibility” [IS02006].

ISO *repeatability* (ISO 3534–2:2006 3.3.5) is defined as precision under “conditions where independent test/measurement results are obtained with the same method on identical test/measurement items in the same test or measuring facility by the same operator using the same equipment within short intervals of time”.

ISO *reproducibility* (ISO 3534–2:2006 3.3.10) is defined as precision under “conditions where independent test/measurement results are obtained with the same method on identical test/measurement items in different test or measurement facilities with different operators using different equipment”

While these definitions are motivated by laboratory use, in a setting where the “test item” is more likely to be a Petri dish culture than a human subject, they still offer a useful sharp definition. In the neuroimaging setting, we find these terms too narrow and unable to capture the range of types of consistency that can be considered. Specifically, not just consistency of the imaging device, or consistency of analysis process, we can consider an expansive concept of consistency, a generalizability of results over different classes of experimental stimuli and context.

In Table A1 we present an incomplete taxonomy of different possible types of consistency, from the ISO repeatability to the widest senses of scientific generalizability. While this taxonomy is useful to see the multifarious nature of “reproducibility”, in the body of this work we use “reproducibility” (without qualification) to mean Roger Peng’s “computational” notion.

**Table B1.Levels of Reproducibility.** .This table provides an incomplete taxonomy of types of consistency of neuroimaging results. For each type of consistency (row), the variable (column) that is held constant (•, bullet) or allowed to vary (D=different) is indicated. In each instance, a bullet (•) indicates the exact same setting; for the variable “Subject” this means the very same acquired data is used, while a double bullet (••) indicates the same subjects are scanned multiple times. Examples of different Experimental Methods include fundamental changes like an event-related design vs. a block design; examples of different Experimental Code/Stimuli, include different sets of pictures used in a visual working memory experiment, or using different paradigm software; an example of different Stimuli Type would be number vs. shape vs. image stimuli in a working memory experiment. An example of different analysis methods for intrasubject (first level) fMRI data would be a confirmatory regression-based modelling vs. an exploratory data-driven method like independent components analysis; examples of different analysis code would be intrasubject fMRI fit with a regression model in two different software packages.

| Type of Stability | Investigator | Experimental Method | Stimuli Type | Experimental Code/Stimuli | Subject Population | Subjects | Scanner | Acquisition Day | Data Analyst | Analysis Method | Analysis Code |
| --- | --- | --- | --- | --- | --- | --- | --- | --- | --- | --- | --- |
| **Measurement Stability** |  |  |  |  |  |  |  |  |  |  |  |
| ISO Repeatability<br>e.g. 30-minute intra-scanner reliability | • | • | • | • | • | •• | • | • | • | • | • |
| ISO Intermediate Reproducibility<br>e.g. 7-day intra-scanner reliability | • | • | • | • | • | •• | • | D | • | • | • |
| ISO Reproducibility<br>e.g. 7-day inter-scanner reliability | • | • | • | • | • | •• | D | D | • | • | • |
| **Analytical Stability** |  |  |  |  |  |  |  |  |  |  |  |
| Analysis Replicability | • | • | • | • | • | • | • | • | • | • | • |
| Collegial Analysis Replicability | • | • | • | • | • | • | • | • | D | • | • |
| Peng's Reproducibility | D | • | • | • | • | • | • | • | D | • | • |
| **Generalisability over ...** |  |  |  |  |  |  |  |  |  |  |  |
| Subjects (Near Replicability) | • | • | • | • | • | D | • | D | • | • | • |
| Labs (Intermediate Replicability) | D | • | • | D | • | D | D | D | D | • | D |
| Experimental & Analytical methods<br>(Far Replicability) | D | D | • | D | • | D | D | D | D | D | D |
| Subject & Stimulus Populations<br>(Hypothesis Generalisability) | D | D | D | D | D | D | D | D | D | D | D |

## Appendix C. Short Descriptions of Fmri Models

While any analysis software consists of myriad modelling decisions, an author must be able to describe the key facets of an analysis in the methods section of their paper. To facilitate this, and to suggest a level of detail that is useful to readers unfamiliar with the software yet not distractingly long, we provide short descriptions for the most commonly used statistical models in widely used software packages.

## C1. Task Fmri

Summaries for AFNI [^44^ http://afni.nimh.nih.gov.] Freesurfer [^45^ https://surfer.nmr.mgh.harvard.edu.], FSL [^46^ http://fsl.fmrib.ox.ac.uk.], & SPM [^47^ http://www.fil.ion.ucl.ac.uk/spm.] are based on versions AFNI_2011_12_21_1014, FreeSurfer 5.3, FSL 5.0.8 and SPM 12 revision 6470, respectively.

**AFNI 1^st^ level – 3dDeconvolve:** Linear regression at each voxel, using ordinary least squares, drift fit with polynomial.

**AFNI 1^st^ level – 3dREMLfit:** Linear regression at each voxel, using generalised least squares with a voxel-wise ARMA(1,1) autocorrelation model, drift fit with polynomial.

**AFNI 2^nd^ level – 3dTtest:** Linear regression at each voxel, using ordinary least squares.

**AFNI 2^nd^ level – 3dMEMA:** Linear mixed effects regression at each voxel, using generalized least squares with a local estimate of random effects variance.

**AFNI 2^nd^ level – 3dMVM:** Multivariate ANOVA or ANCOVA at each voxel.

**AFNI 2^nd^ level – 3dLME:** General linear mixed-effects modeling at each voxel, with separate specification of fixed and random variables.

**Freesurfer 1st Level – selxavg3-sess:** Linear regression at each surface element, using generalized least squares with a element-wise AR(1) autocorrelation model, drift fit with polynomial.

**Freesurfer 2st Level – mri_glmfit:** Linear regression at each surface element, using ordinary least squares.

**FSL 1^st^ level:** Linear regression at each voxel, using generalized least squares with a voxel-wise, temporally and spatially regularized autocorrelation model, drift fit with Gaussian-weighted running line smoother (100s FWHM).

**FSL 2^nd^ level – “OLS”:** Linear regression at each voxel, using ordinary least squares.

**FSL 2^nd^ level – “FLAME1”:** Linear mixed effects regression at each voxel, using generalized least squares with a local estimate of random effects variance.

**SPM 1^st^ level:** Linear regression at each voxel, using generalized least squares with a global approximate AR(1) autocorrelation model, drift fit with Discrete Cosine Transform basis (128s cut-off).

**SPM 2^nd^ level – no repeated measures:** Linear regression at each voxel, using ordinary least squares.

**SPM 2^nd^ level – repeated measures:** Linear regression at each voxel, using generalized least squares with a global repeated measures correlation model.

## C2. Single-Modality ICA

Methods for ICA analyses are not as consolidated as mass univariate linear modelling, but we provide short summaries of some typical analyses in GIFT [^48^ http://mialab.mrn.org/software/gift.] and MELODIC [^49^ http://fsl.fmrib.ox.ac.uk/fsl/fslwiki/MELODIC.] (alphabetical order), based on versions GIFTv3.0a and FSL 5.0.8, respectively. [Optional aspects, depending on particular variants used, indicated in brackets.]

**GIFT, single-subject fMRI with ICASSO stability:** Spatial ICA estimated with infomax where scaling of original data, spatial components and time courses constrained to unit norm, resulting best-run selected from 10 runs; post-ICA Z statistics produced for maps, between temporal component correlation (Functional Network Correlation), time courses, spectra, tested within a GLM framework.

**GIFT, multi-subject PCA-based back-reconstruction with ICASSO stability:** Single-subject PCA followed by temporal concatenation, group-level PCA and then spatial ICA with infomax; calculation of single subject maps using PCA-based back-reconstruction, resulting best-run selected from 10 runs; post-ICA Z statistics produced for maps, time courses, spectra, and between temporal component correlation (Functional Network Correlation) tested within a GLM framework. [Time-varying states computed using moving window between temporal components (Dynamic Functional Network Correlation).]

**GIFT, spatio-temporal (dual) regression of new data:** Using provided component maps calculates per-subject components from new data using regression-based back-reconstruction; produces component maps, time courses and spectra and between temporal component correlation (Functional Network Correlation) tested within a GLM framework.

**GIFT, spatial ICA with reference:** Spatial ICA using one or more provided seed or component maps. Components found by joint maximization of non-Gaussianity and similarity to spatial maps resulting in subject specific component maps and timecourses for each subject, scaled to Z-scores, following by testing voxelwise (within network connectivity), between temporal component correlation (Functional Network Correlation), spectra, tested within a GLM framework.

**GIFT, source based morphometry of gray matter maps:** Spatial ICA of multi-subject gray matter segmentation maps (from SPM, FSL, etc) resulting in spatial components and subject-loading parameters tested within a GLM framework.

**MELODIC, single-subject ICA:** Spatial ICA estimated by maximising non-Gaussian sources, using robust voxel-wise variance-normalisation of data, automatic model-order selection and Gaussian/Gamma mixture-model based inference on component maps.

**MELODIC, group level (concat ICA):** Temporally concatenation of fMRI data, followed by spatial ICA estimated by maximising non-Gaussian sources, using using robust voxel-wise variance-normalisation of data, automatic model order selection and Gaussian/Gamma mixture-model based inference on component maps

**MELODIC, group-level (tensor-ICA):** Higher-dimensional decomposition of all fMRI data sets into spatial, temporal and subject modes; automatic model order selection and Gaussian/Gamma mixture-model based inference on component maps

**MELODIC dual regression:** Estimation of subject-specific temporal and spatial modes from group-level ICA maps or template maps using spatial followed by temporal regression.

## C3. Multi-Modalitiy ICA

Available multi-modality ICA methods include FIT [^50^ http://mialab.mrn.org/software/fit.] and FSL-FLICA [^51^ http://fsl.fmrib.ox.ac.uk/fsl/fslwiki/FLICA.] (alphabetical order), based on versions FITv2.0c and flica_2013–01–15, respectively.

**FIT, joint ICA, two-group, fMRI + EEG fusion:** Joint spatial ICA of GLM contrast maps and temporal ICA of single or multi-electrode event-related potential time course data (can be non-concurrent) with infomax ICA; produces joint component maps (each with an fMRI component map and ERP component timecourse(s)) and subject loading parameters which are then tested for group differences with a GLM framework.

**FIT, N-way fusion using multiset CCA+joint ICA:** Multiset canonical correlation analysis applied to several spatial maps to extract components, then submitted to spatial ICA with infomax ICA; produces multi-modal component maps and subject-specific loading parameters which are tested within a GLM framework.

**FIT, parallel ICA, fusion of gray matter maps and genetic polymorphism array data:** Joint spatial ICA of gray matter segmentation maps and genetic ICA of single nucleotide polymorphism data performed through a maximization of independence among gray matter components, genetic components, and subject-wise correlation among one or more gray matter and genetic components. Produces linked and unlinked gray matter and genetic components and subject loading parameters which are then tested within a GLM framework.

**FSL-FLICA multi-subject/multi-modality (Linked-ICA):** ICA-based estimation of common components across multiple image modalities, linked through a shared subject-courses.

## Appendix D. Itemized Lists of best Practices and Reporting Items

This section contains checklists for practices and items to report. Each item has been included because it is an essential piece of information needed to understand, evaluate and reproduce an experiment. Authors should strive to include all these items, but items marked as “Mandatory” are particularly crucial, and a published work cannot be considered complete without such information.

*The rest of this page intentionally left blank*.

**Table D.1.**
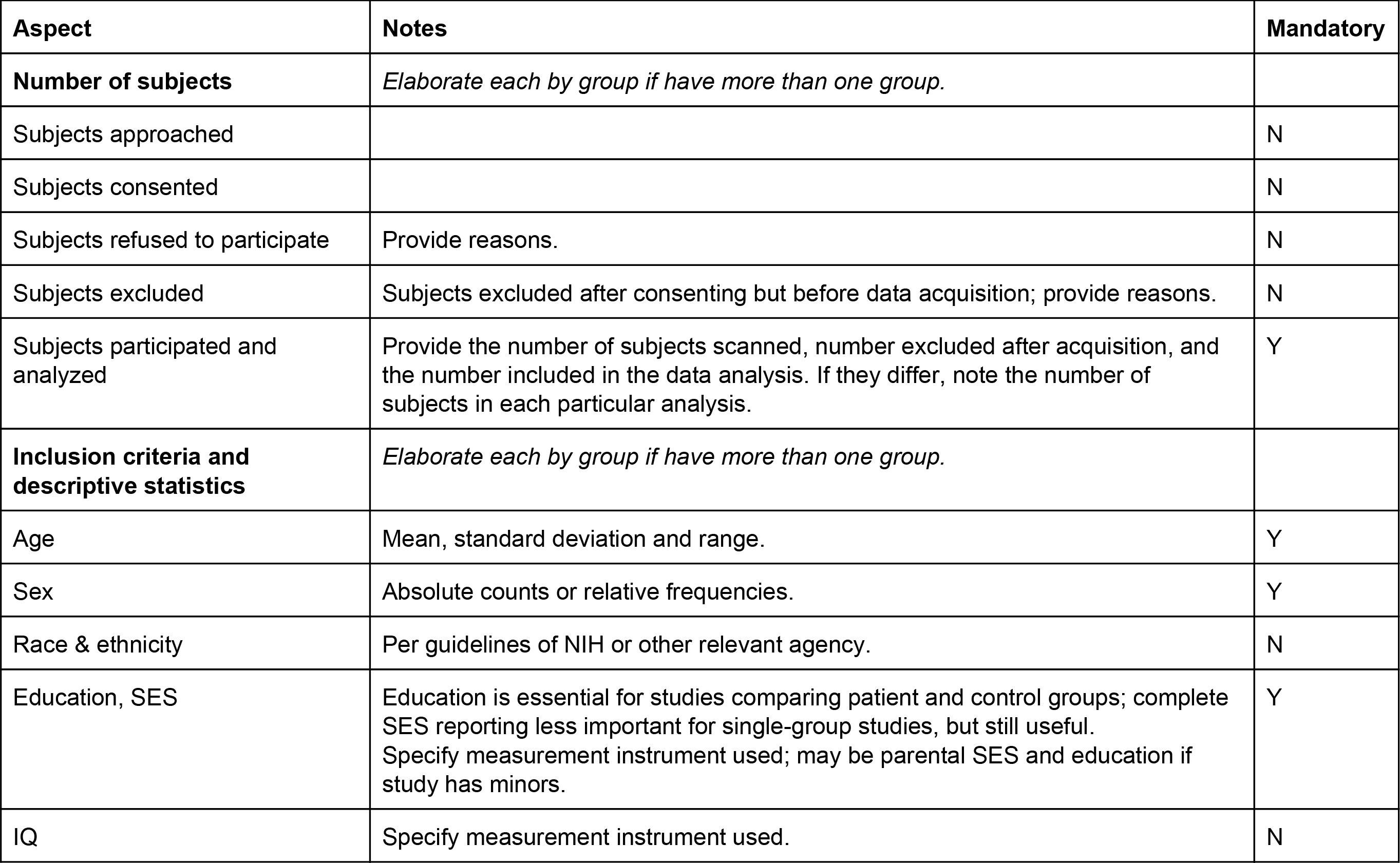
Experimental Design Reporting

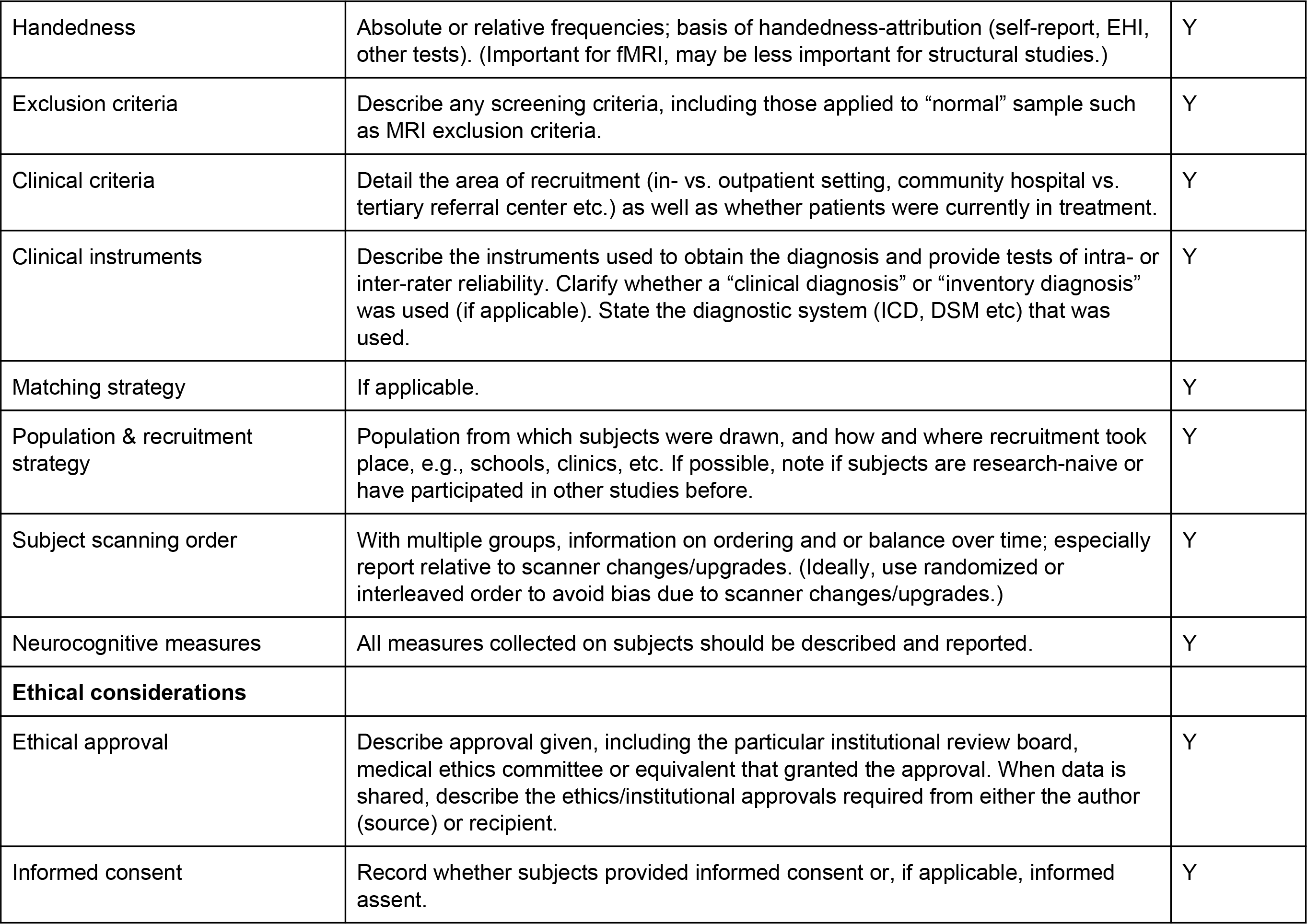

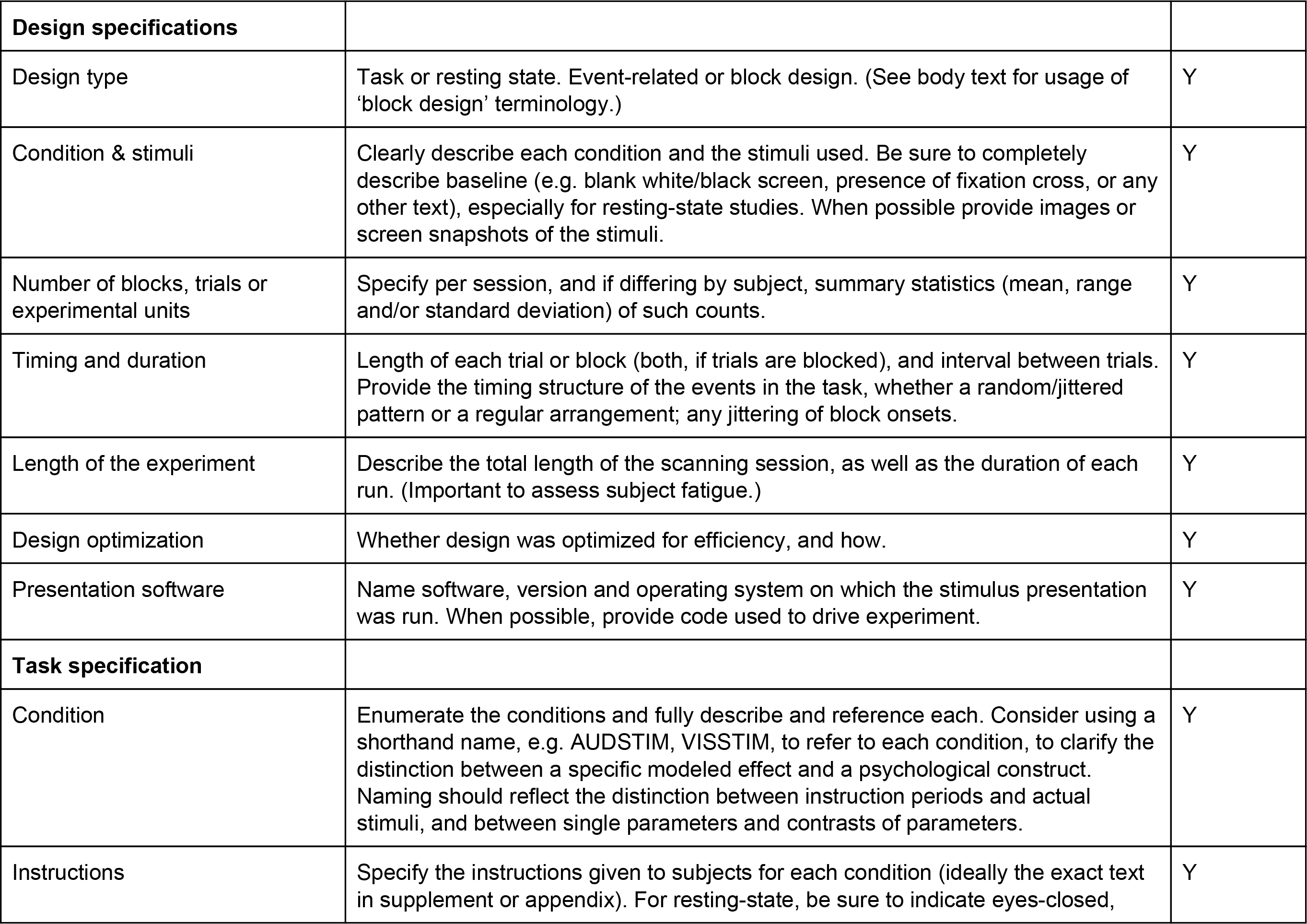

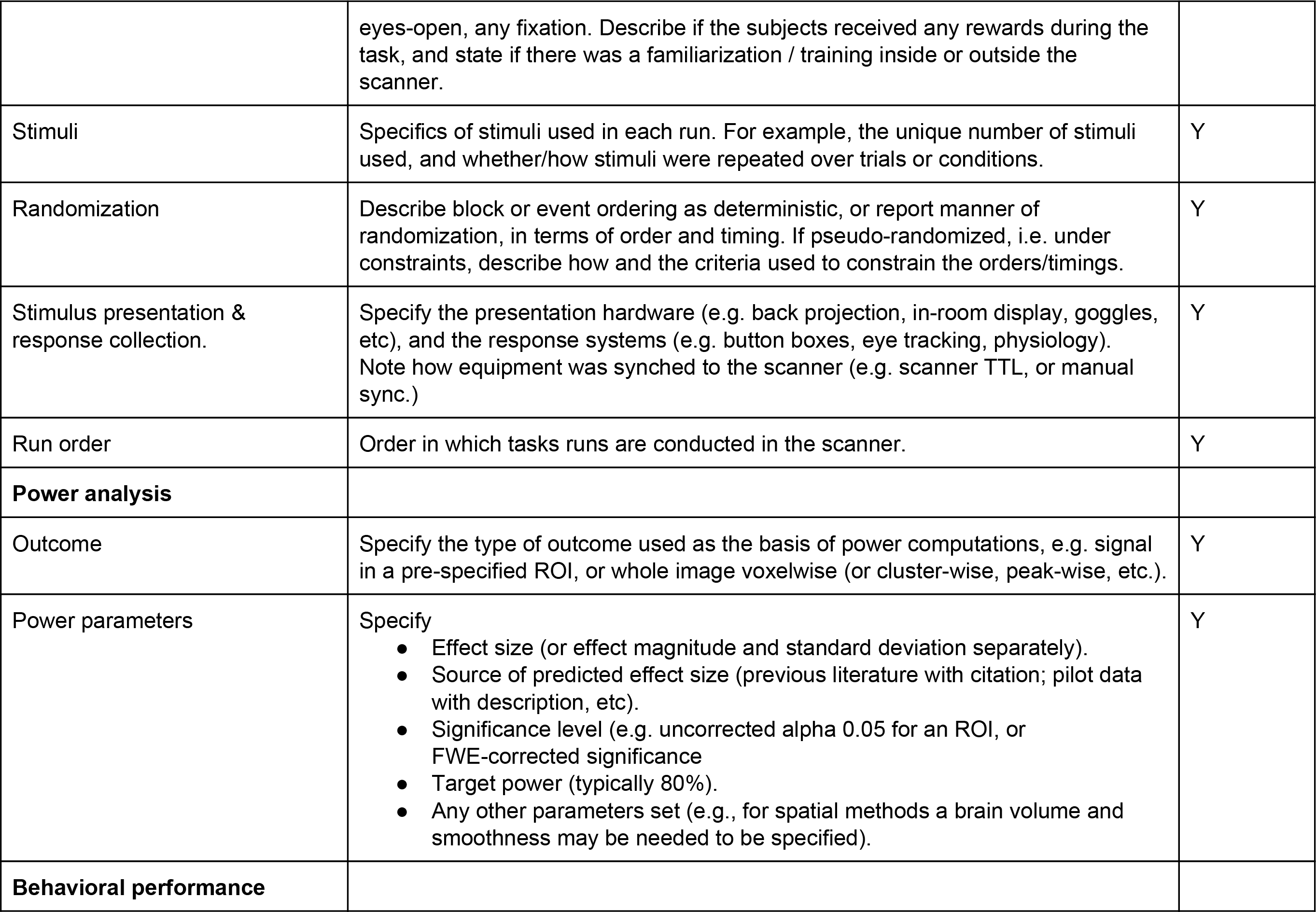

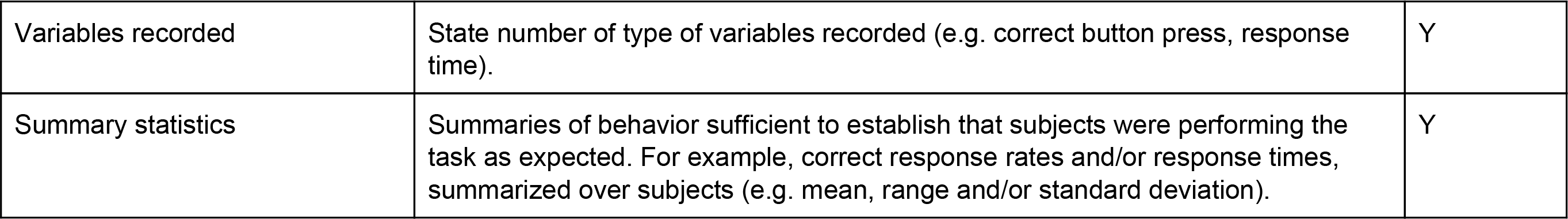

**Table D.2.**
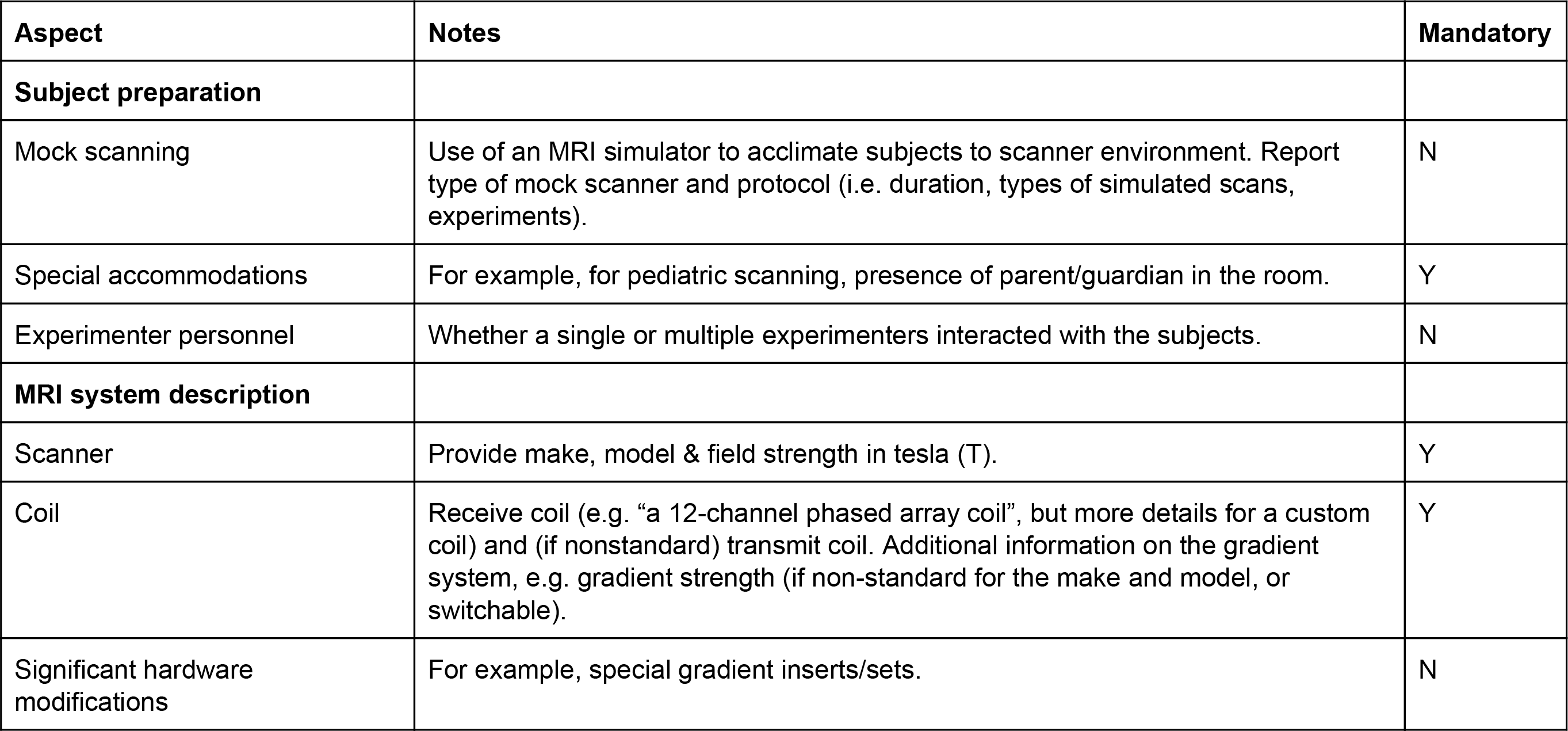
Acquisition Reporting

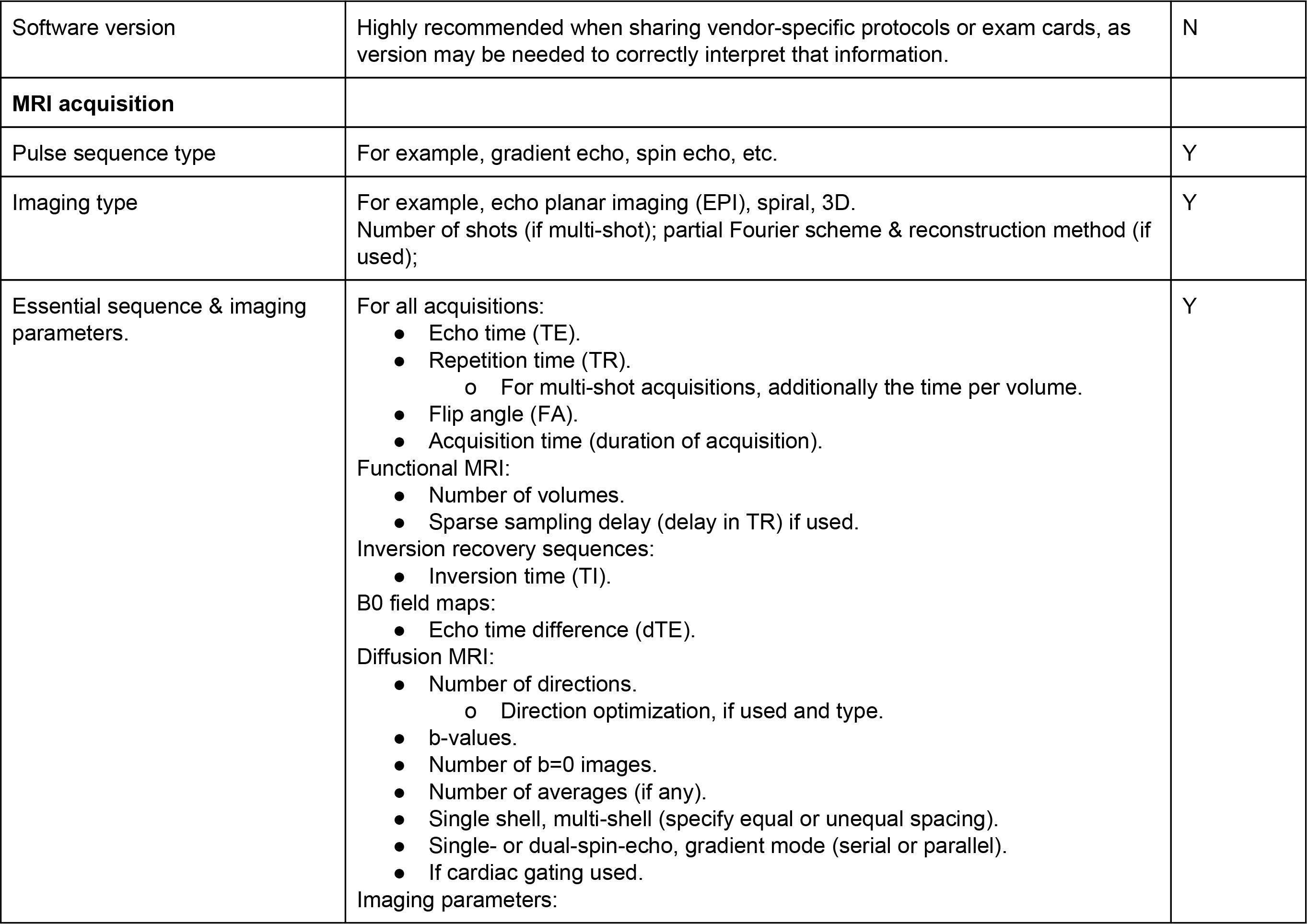

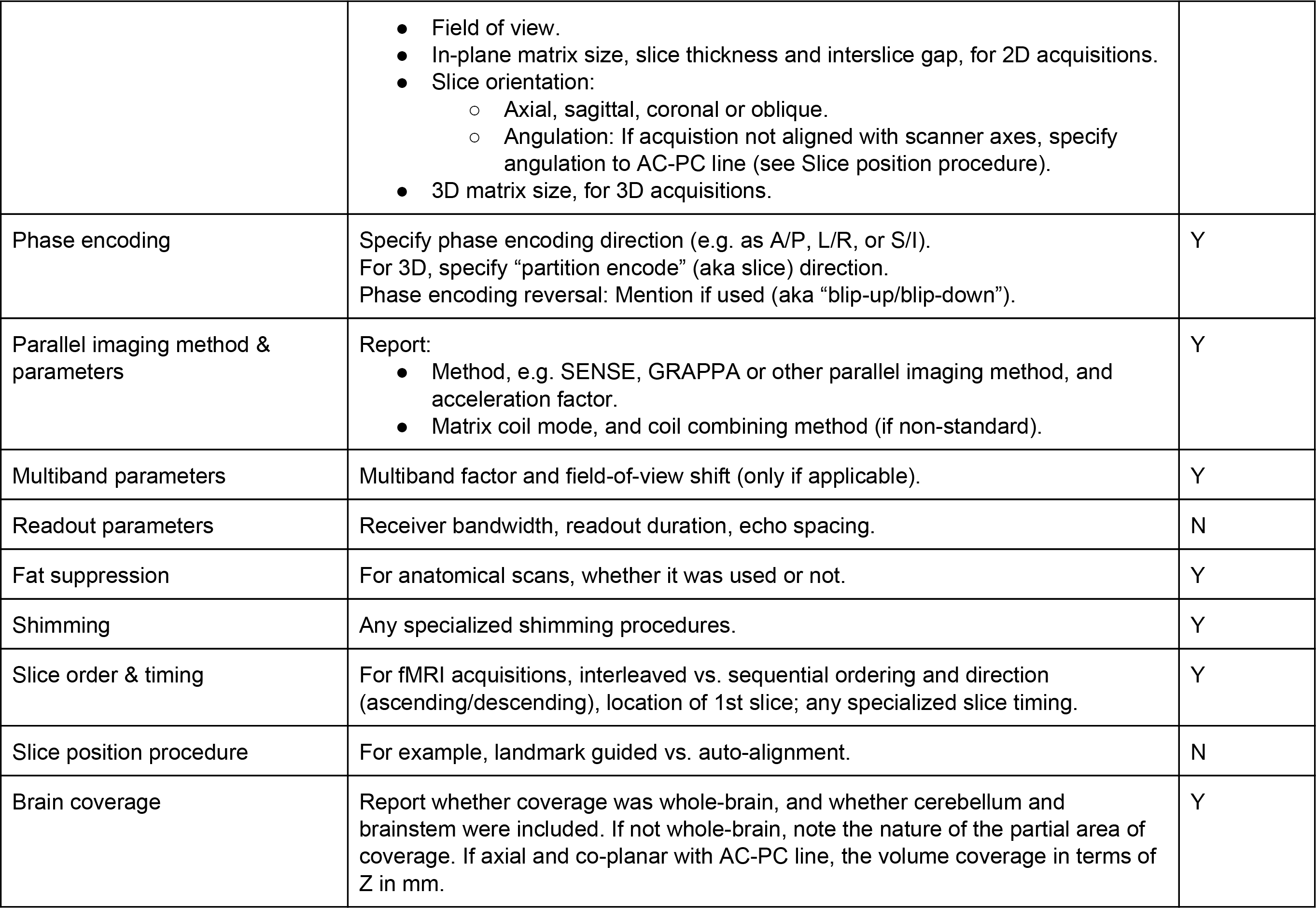

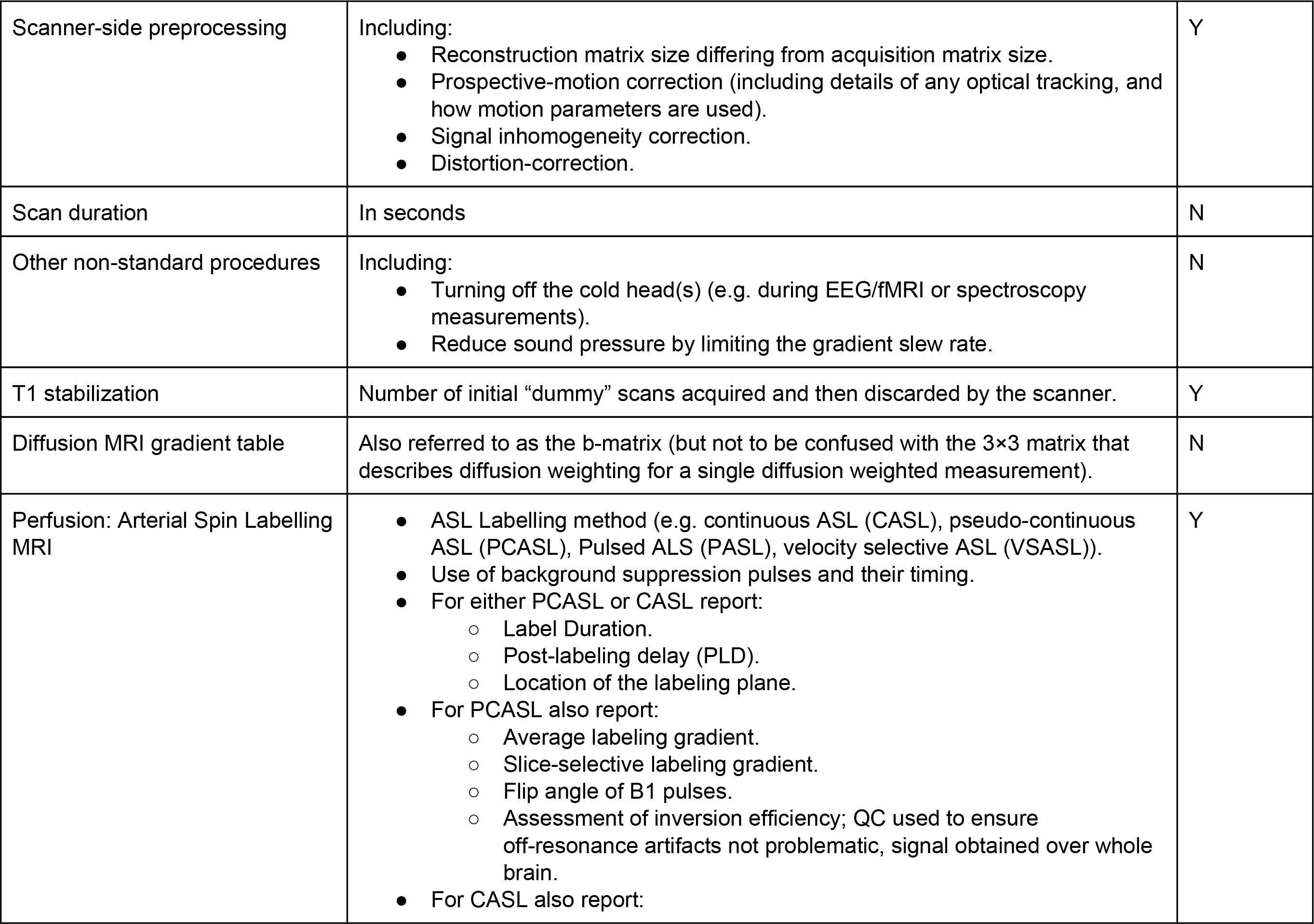

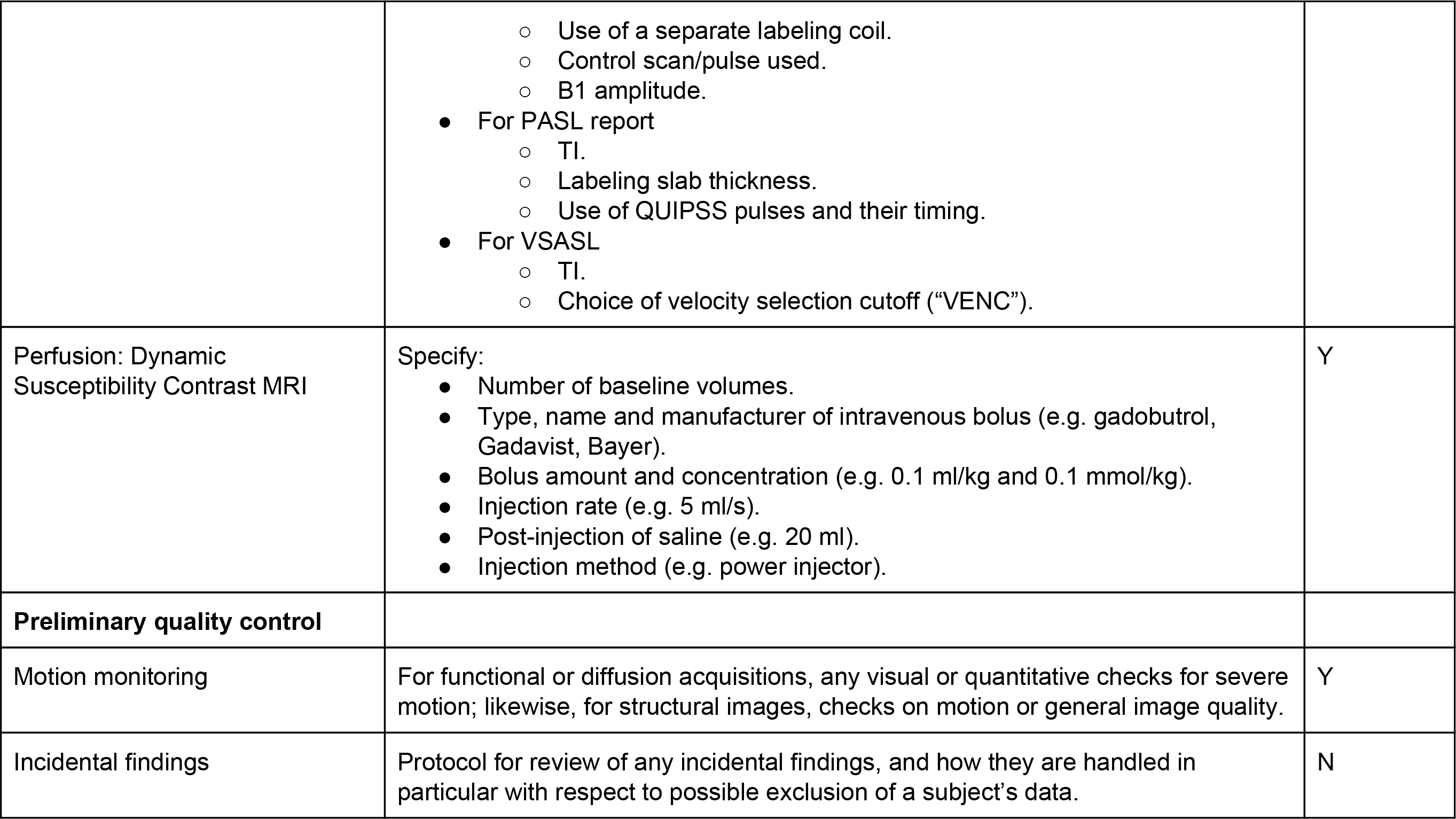

**Table D.3.**
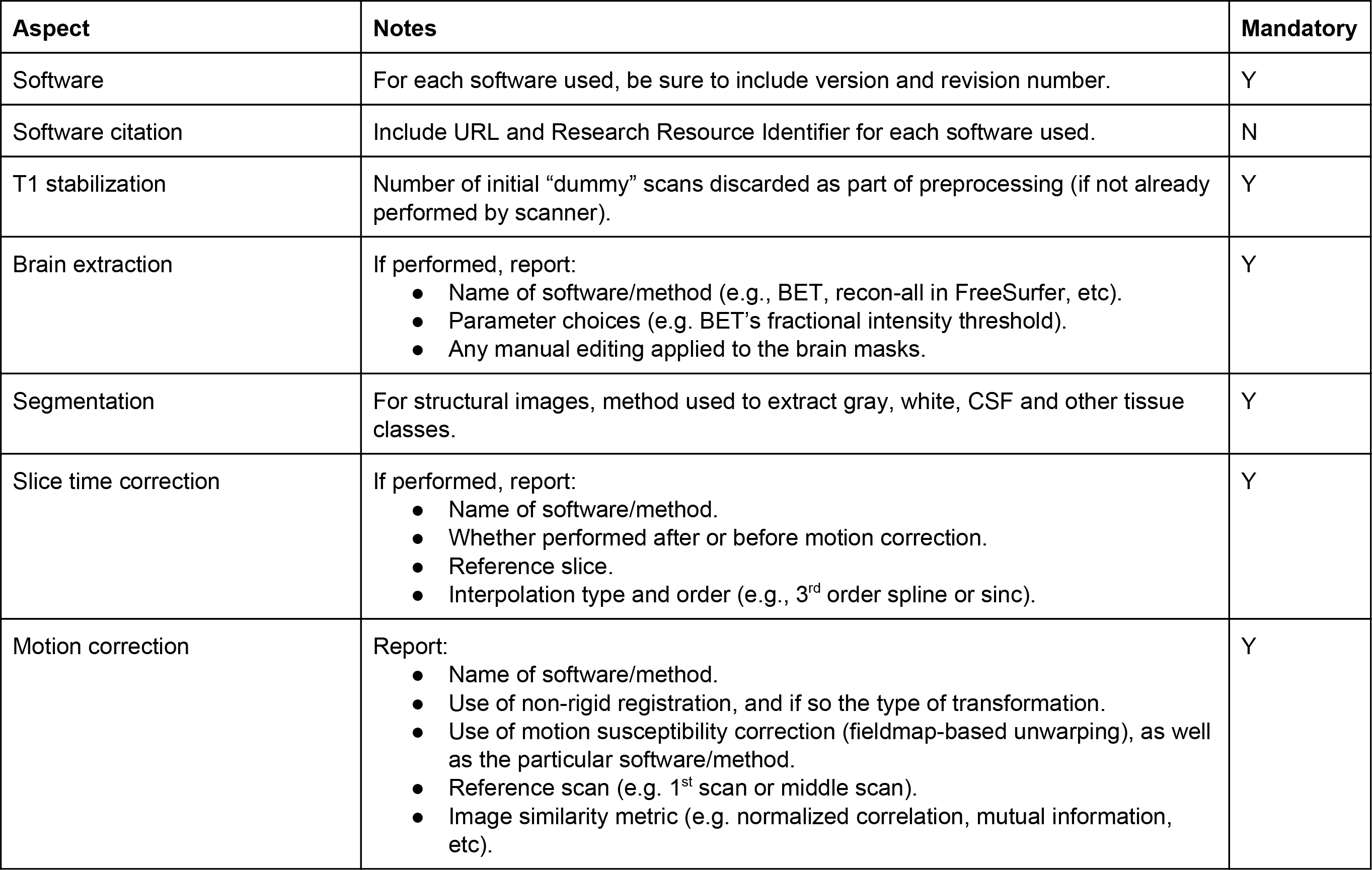
Preprocessing Reporting

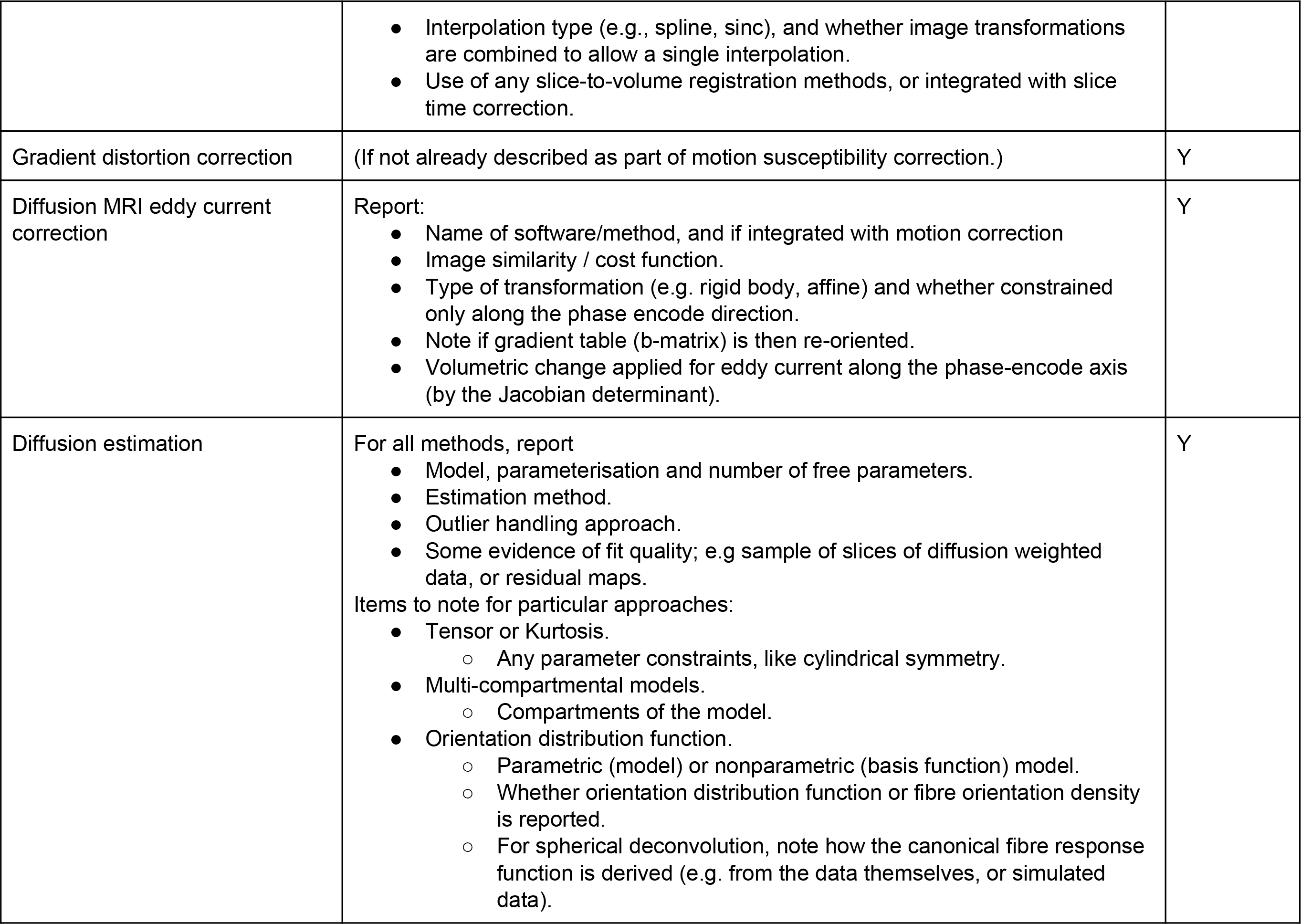

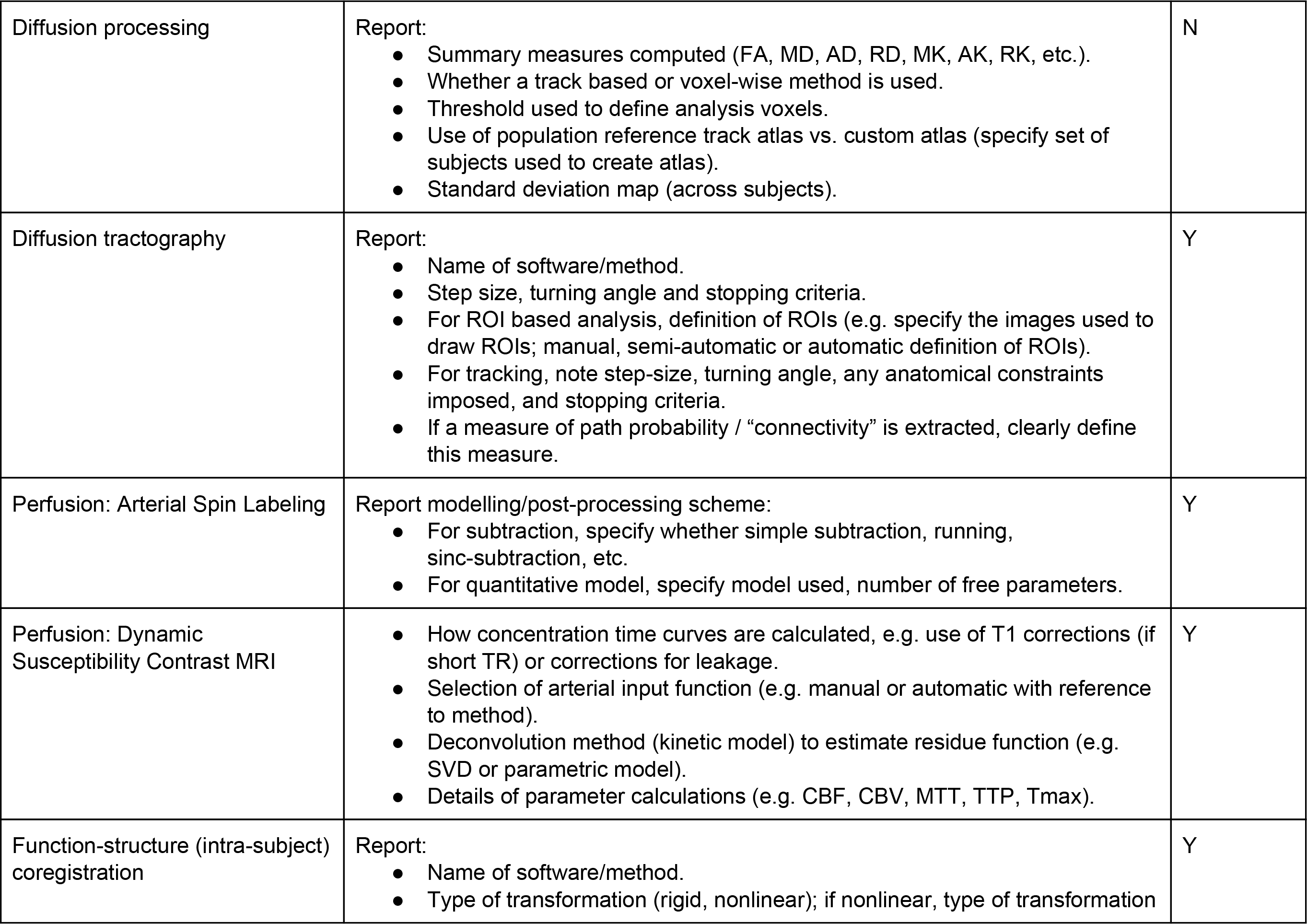

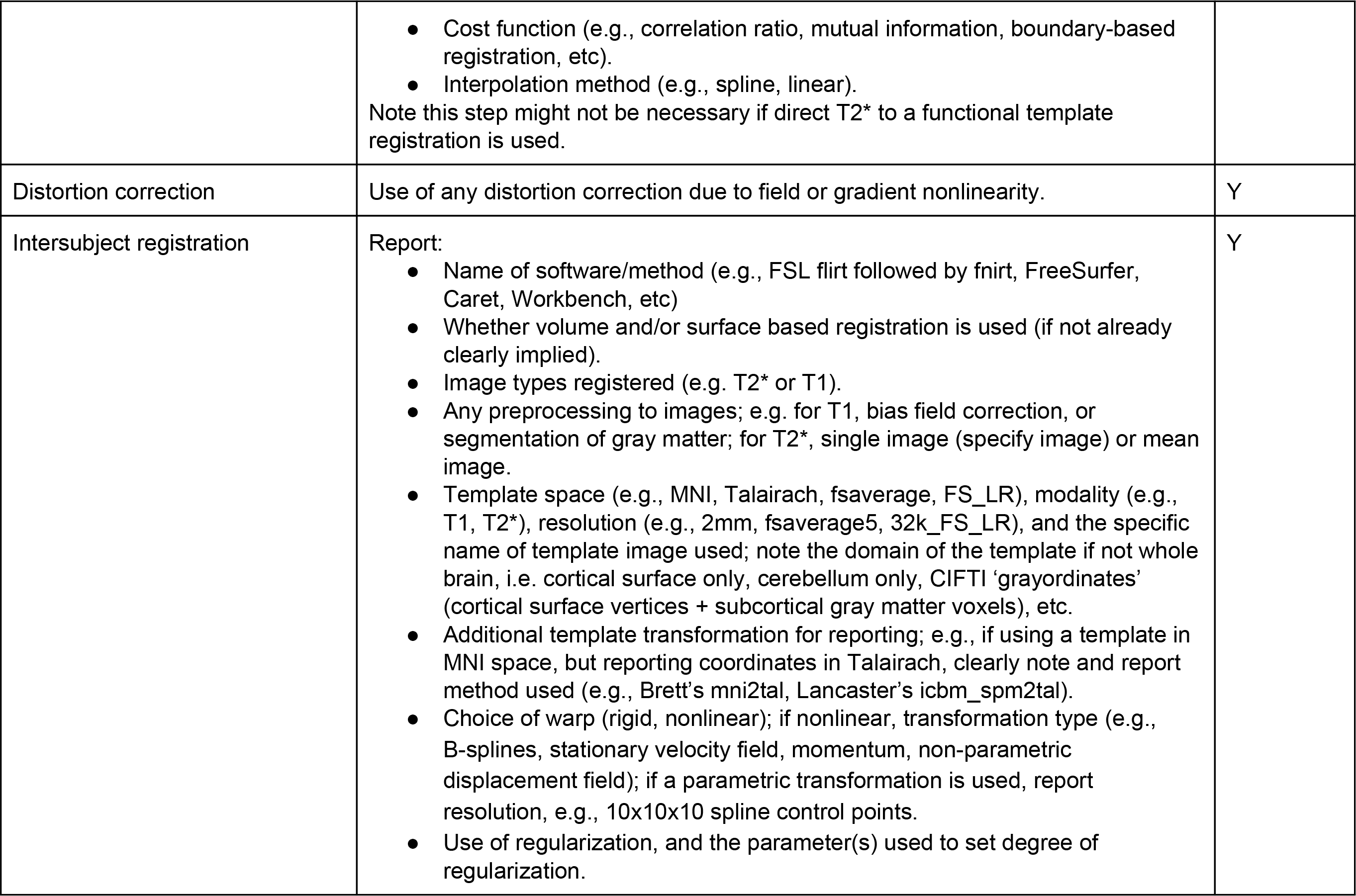

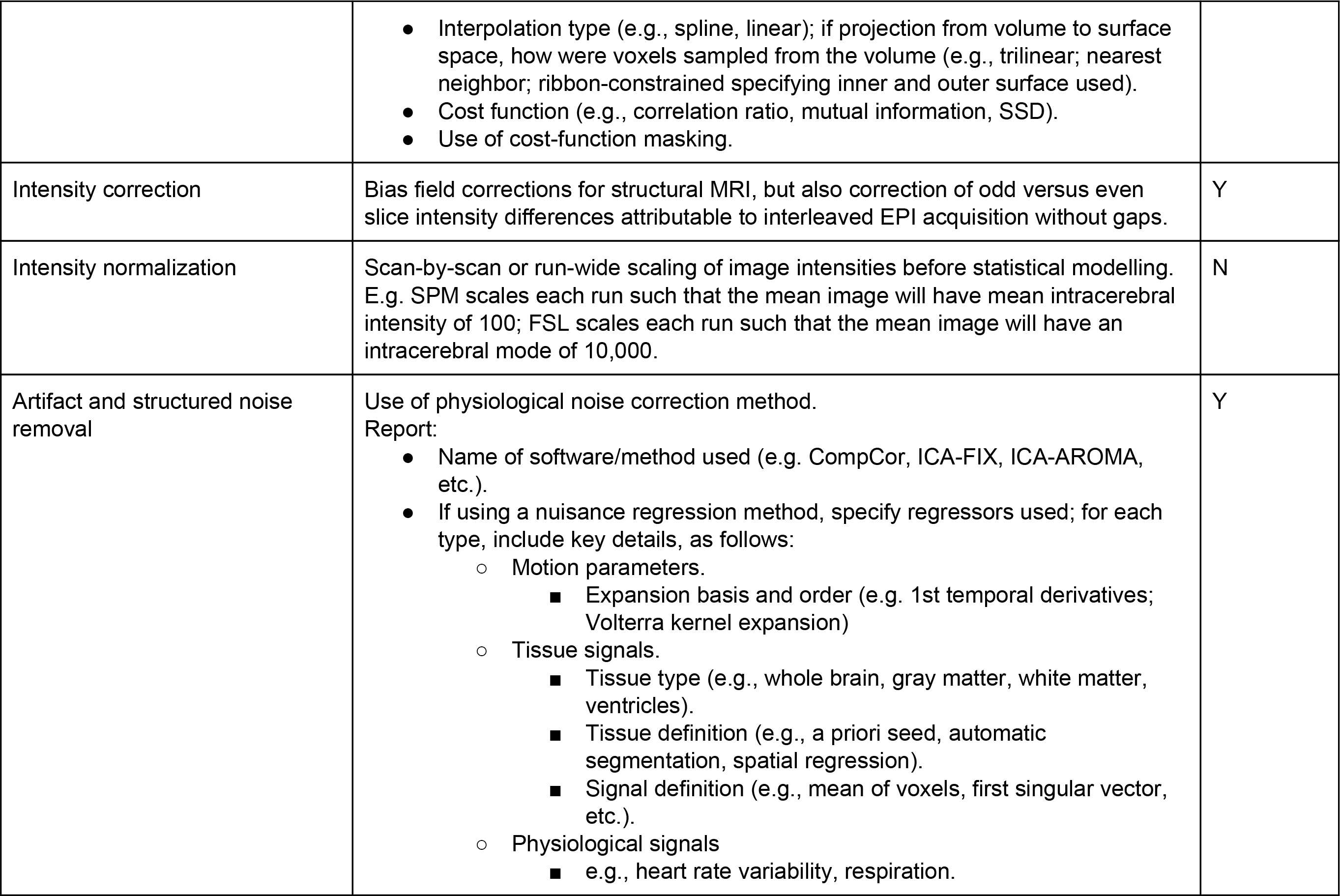

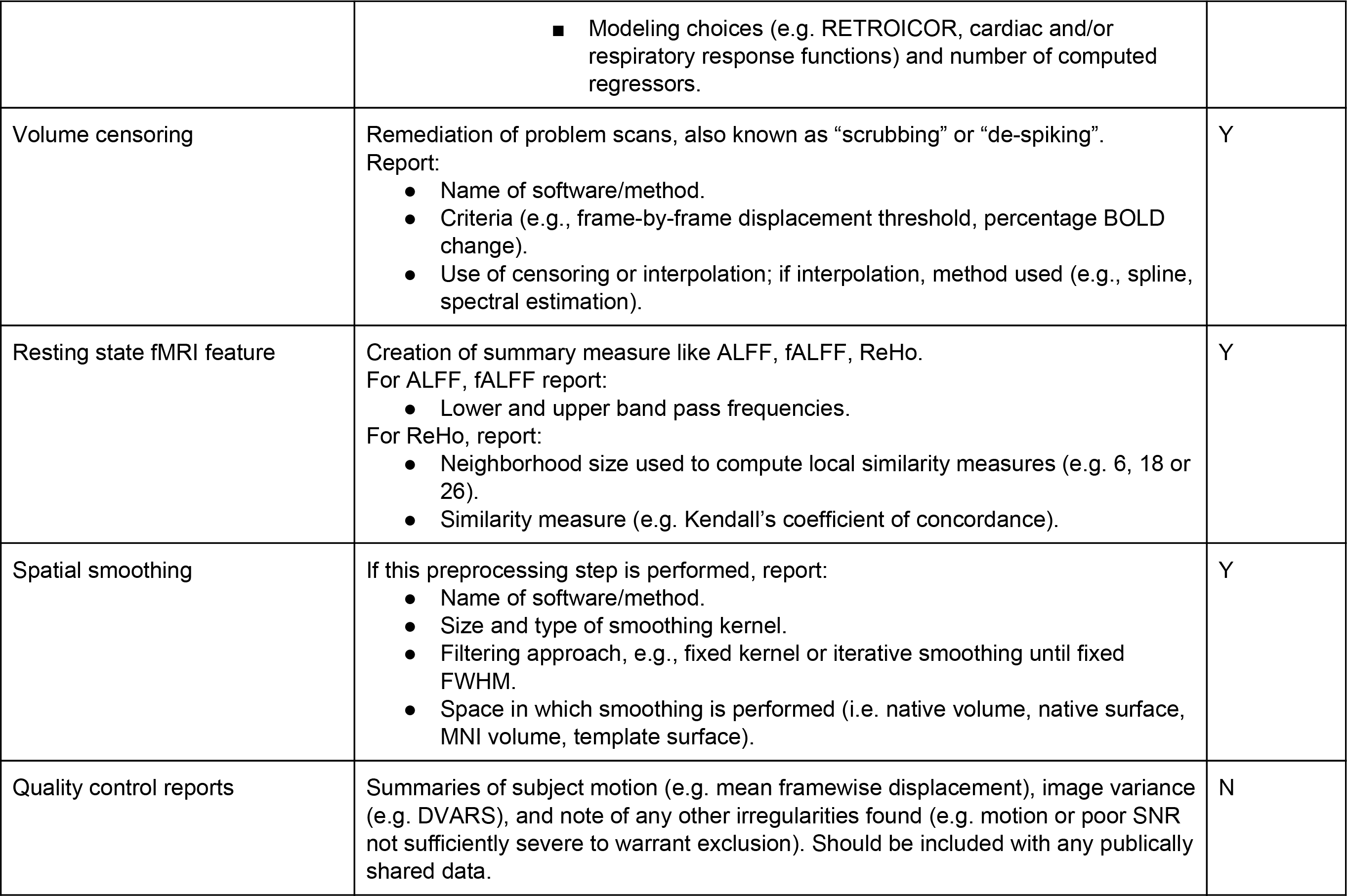

**Table D.4.**
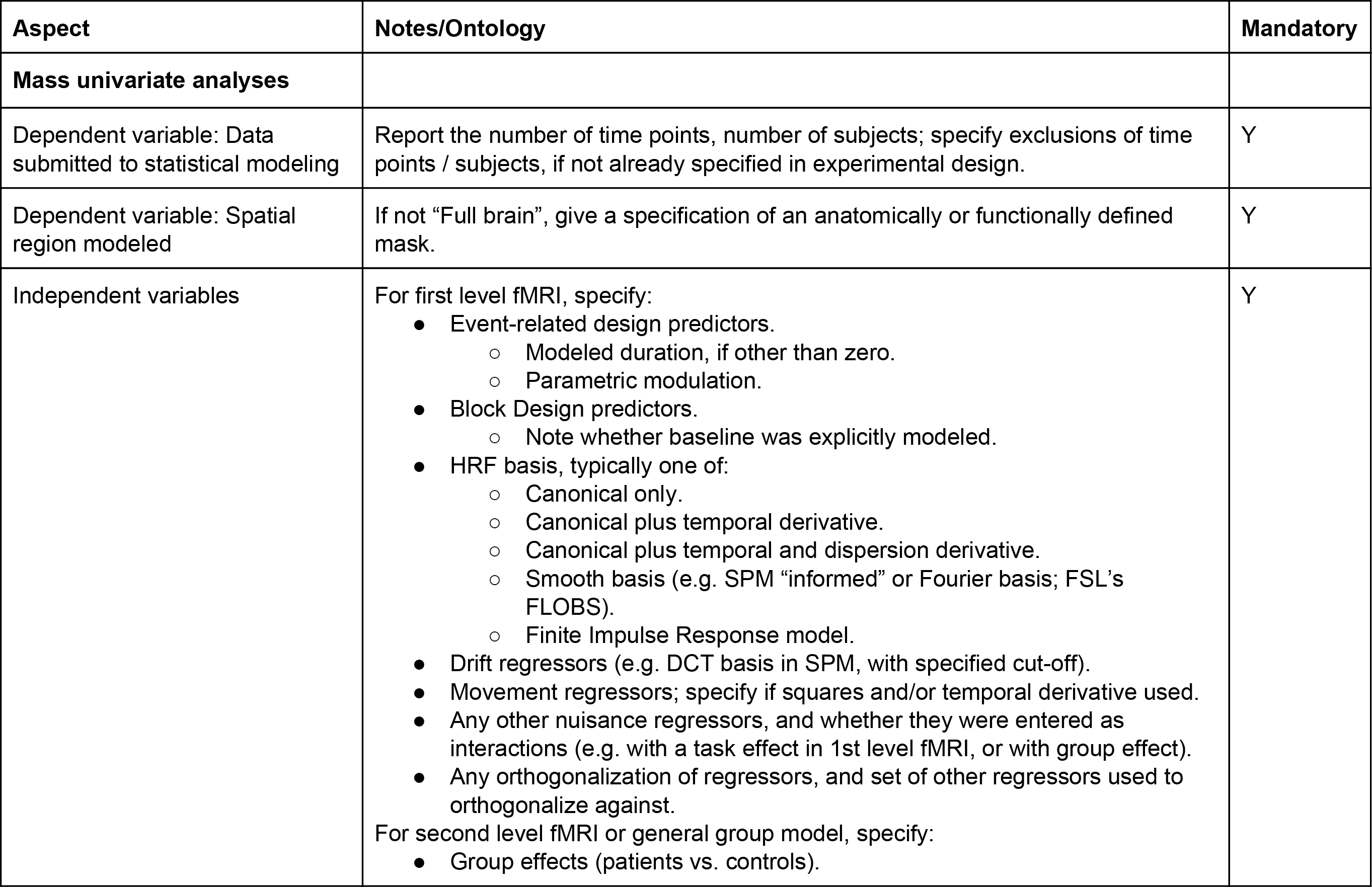
Statistical Modeling & Inference

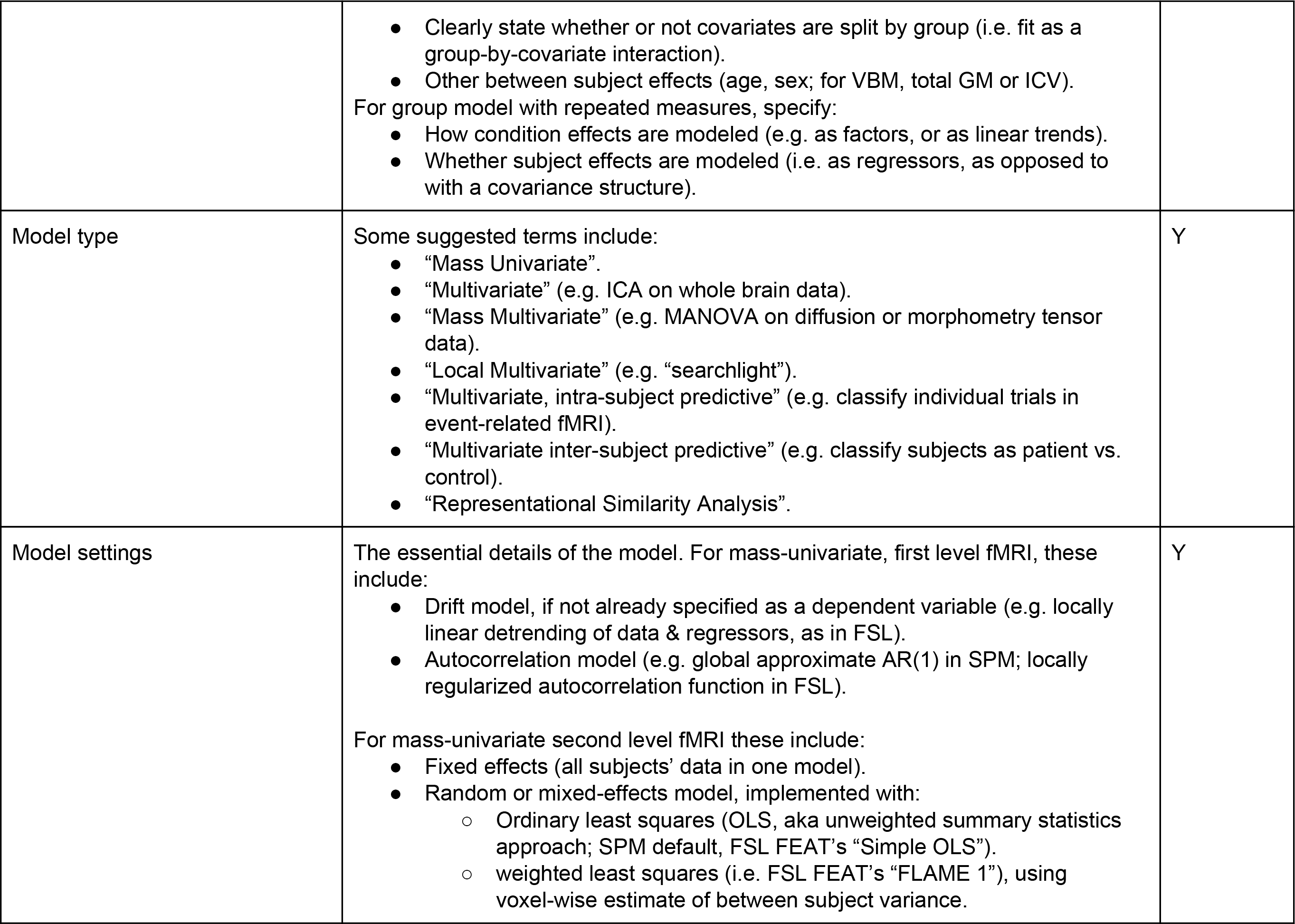

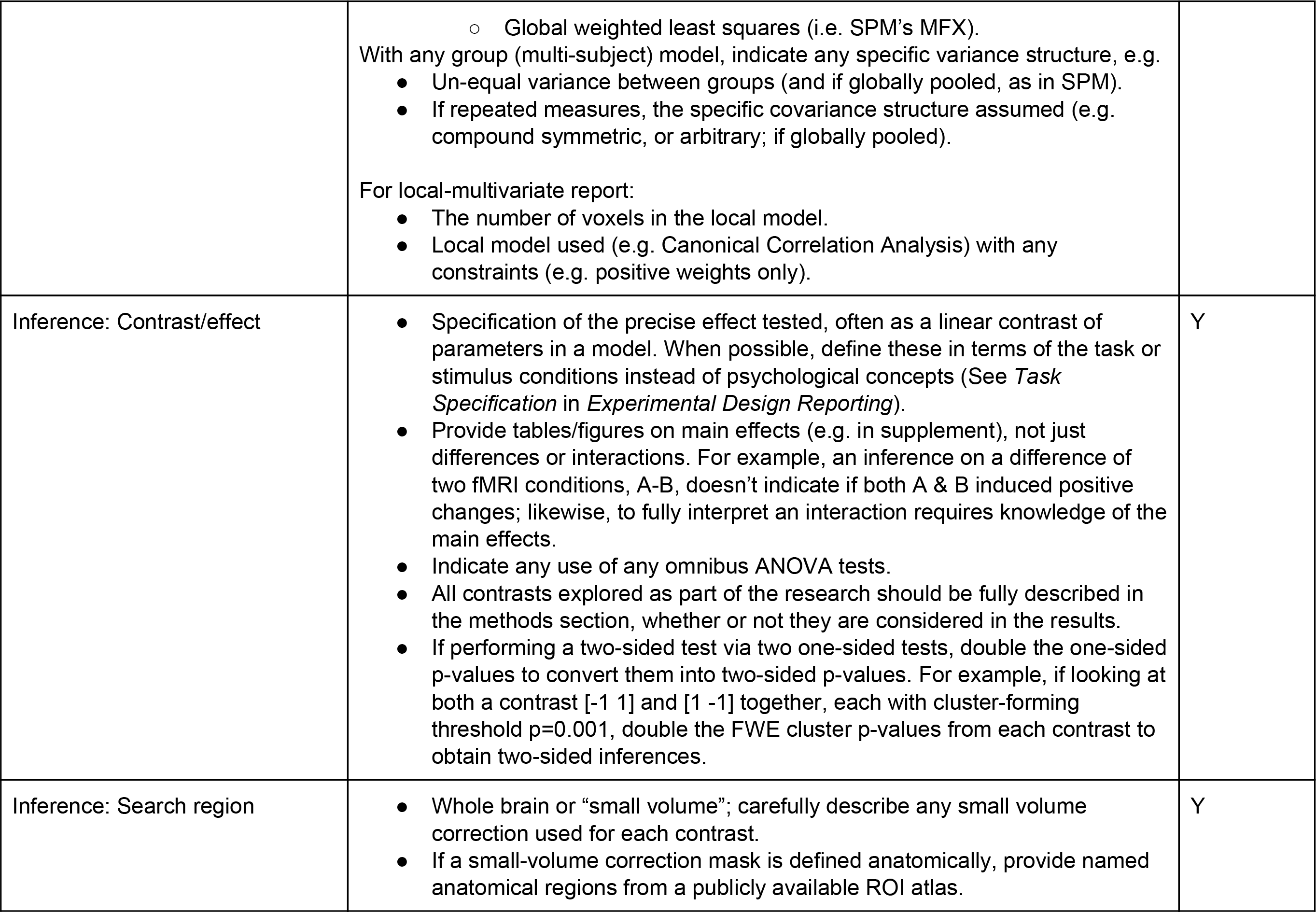

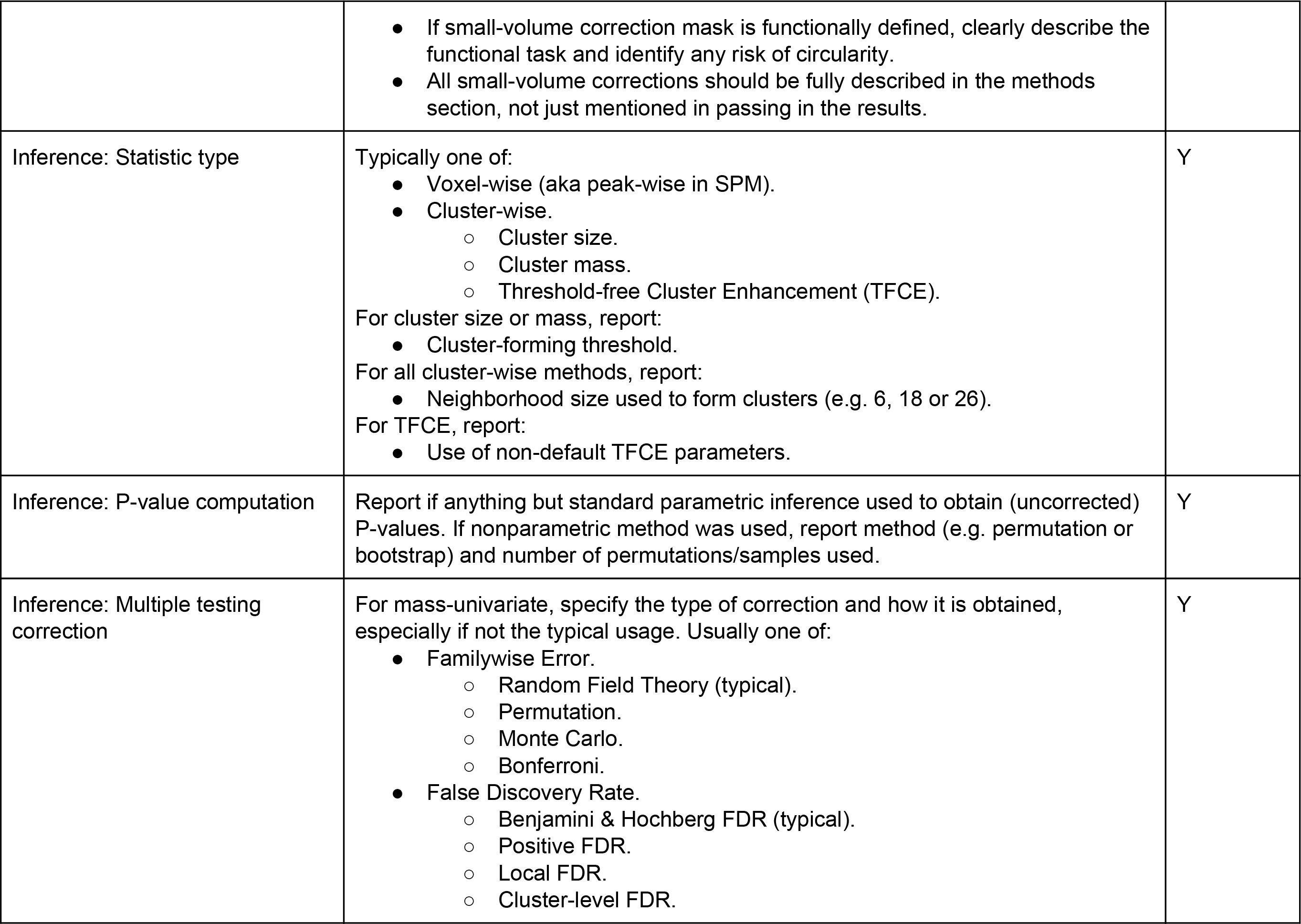

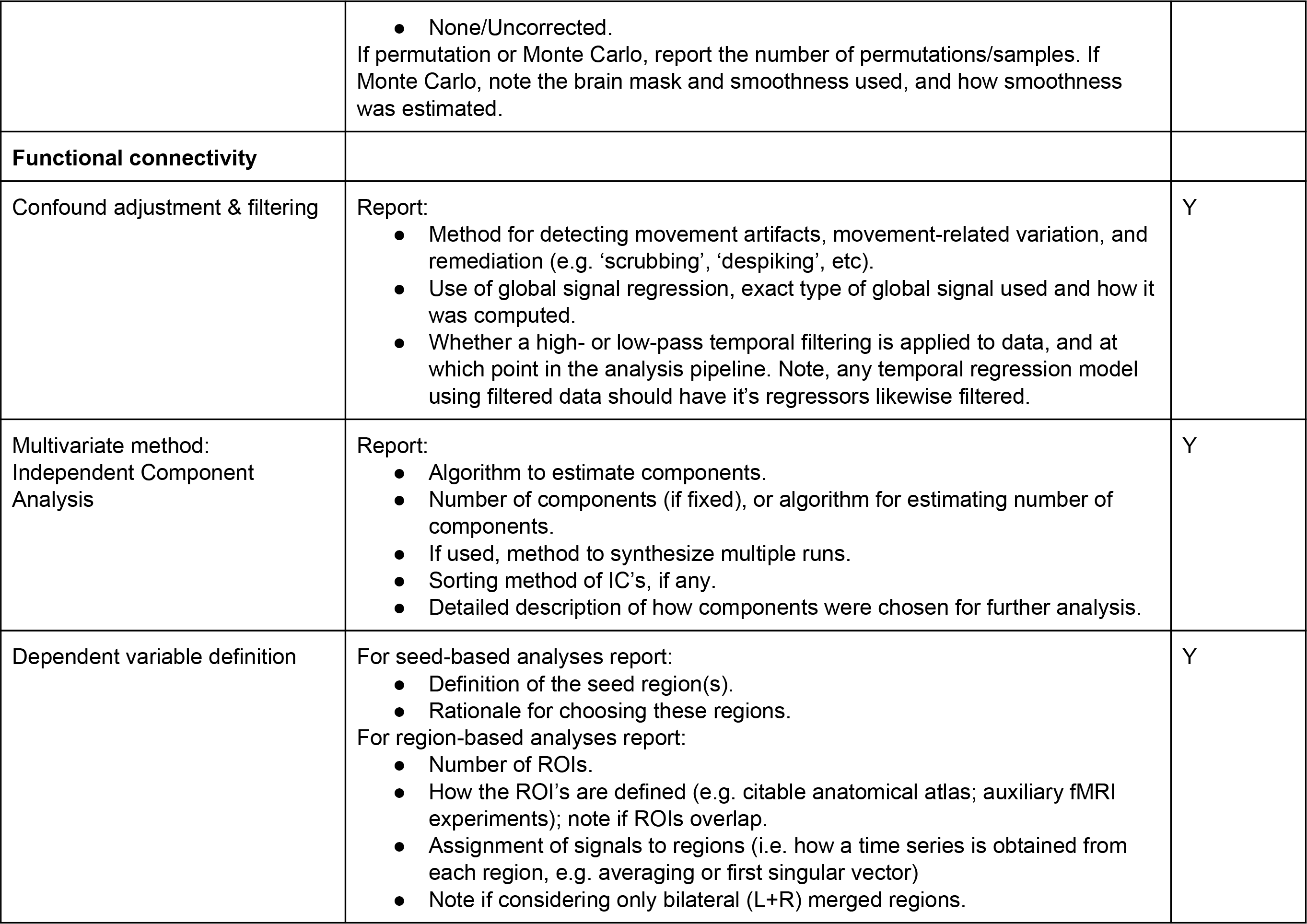

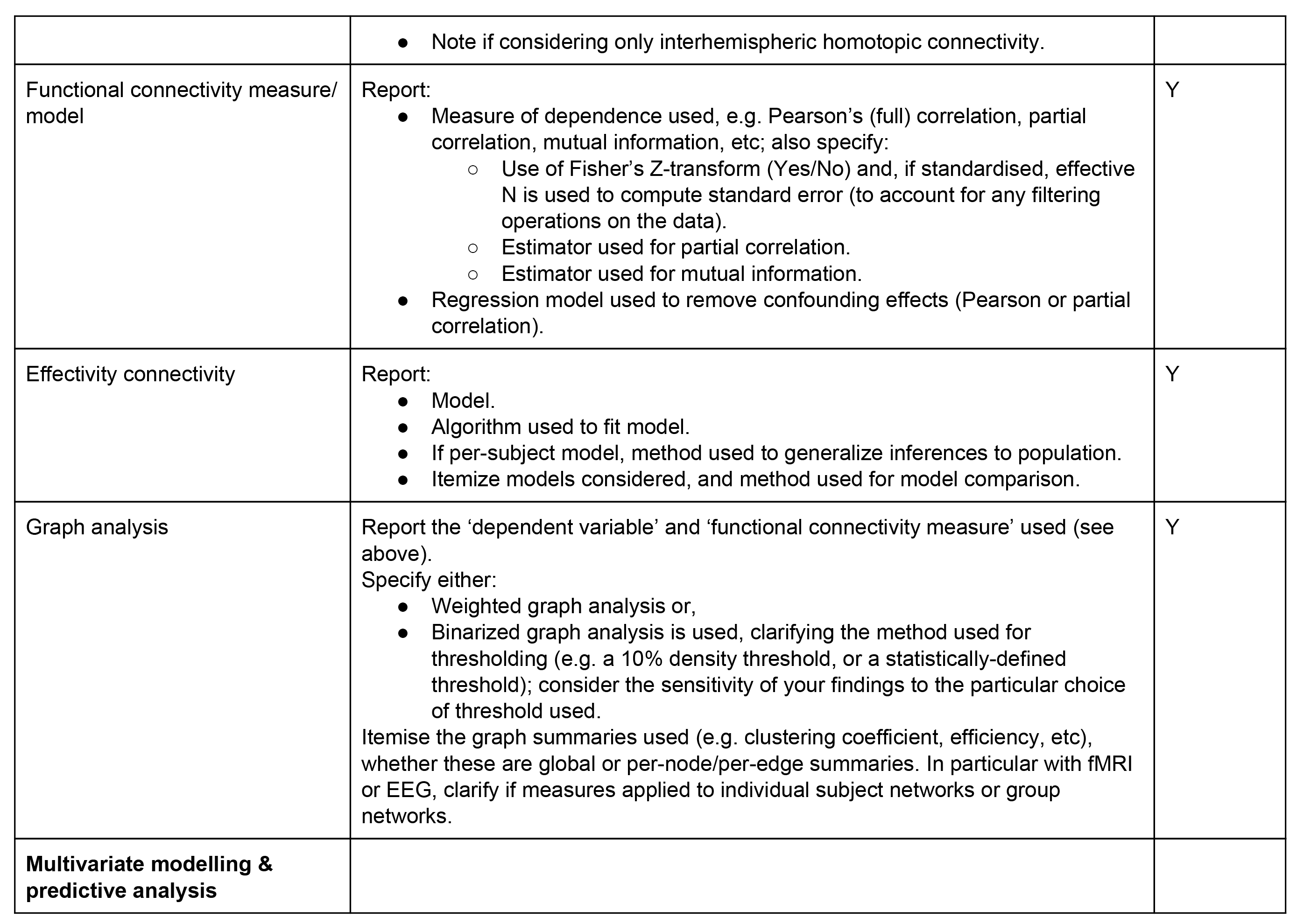

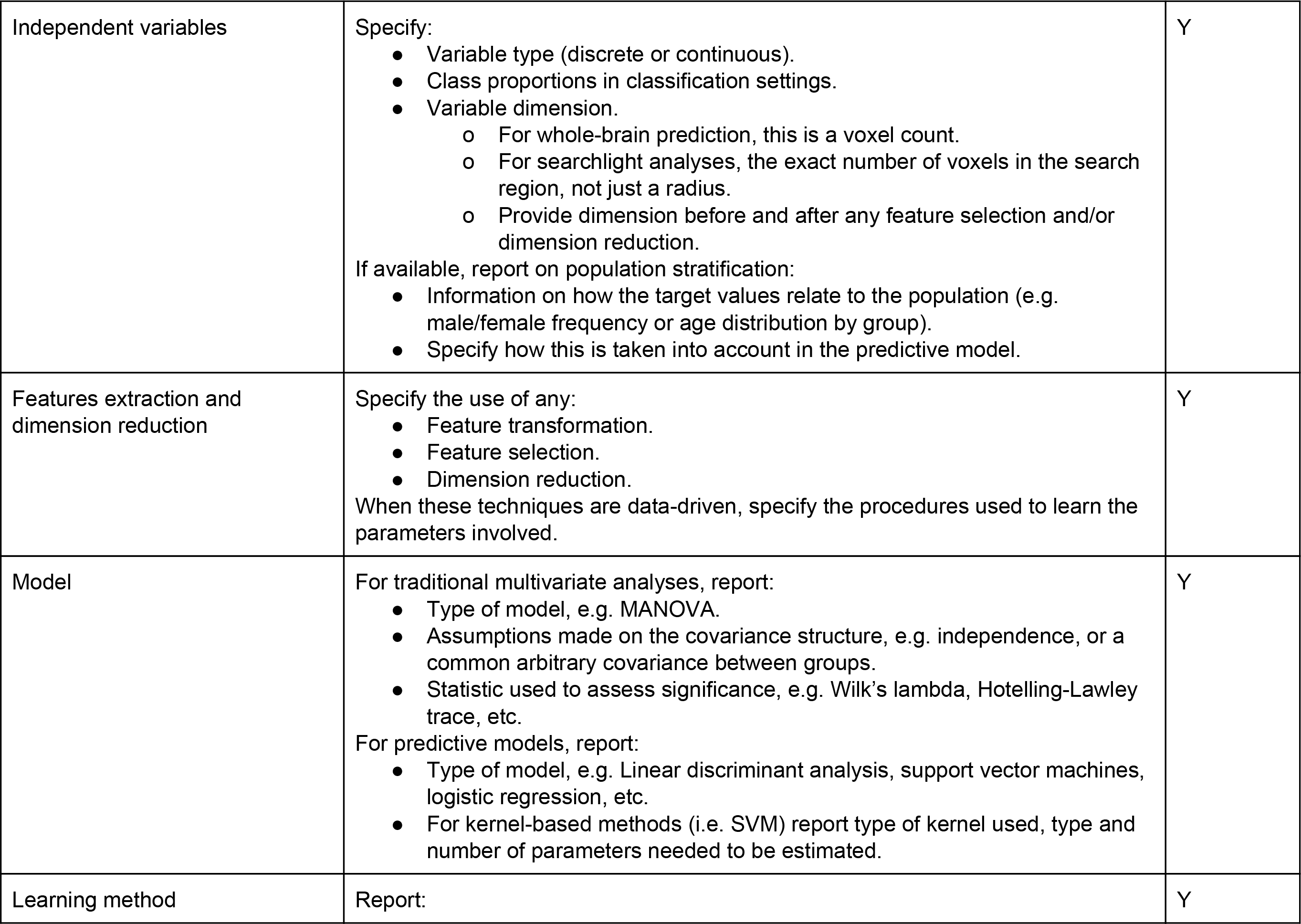

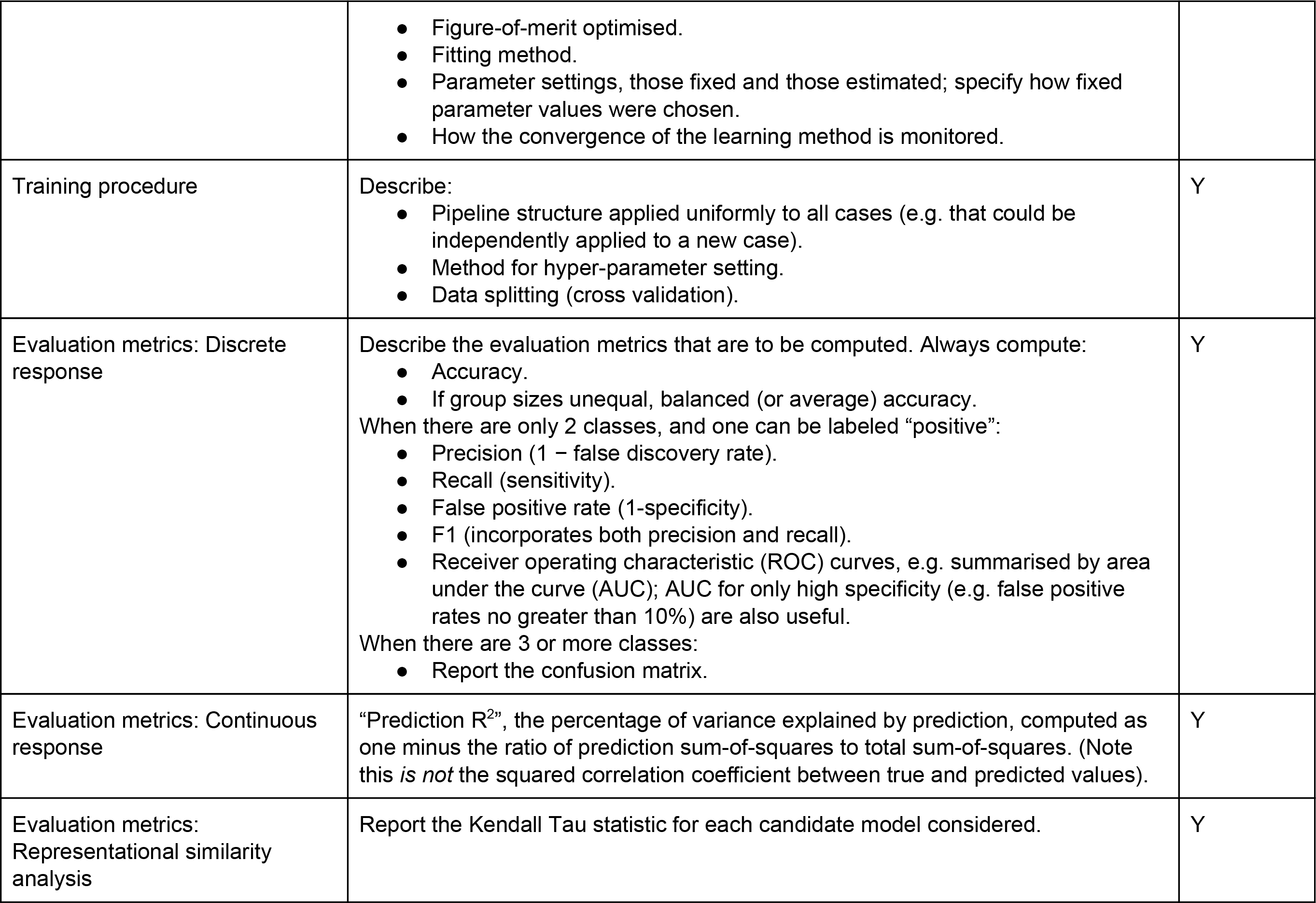

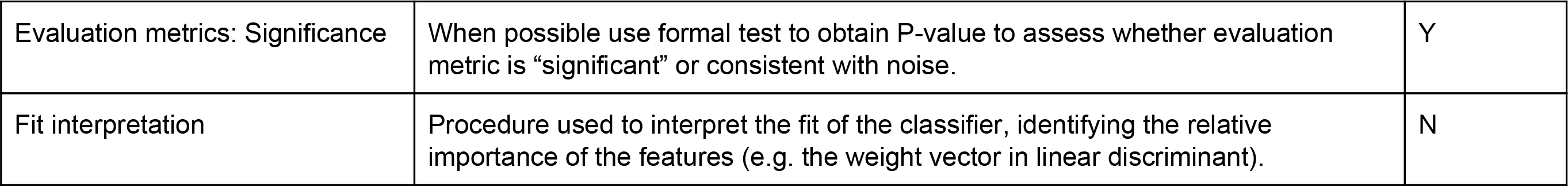

**Table D.5.**
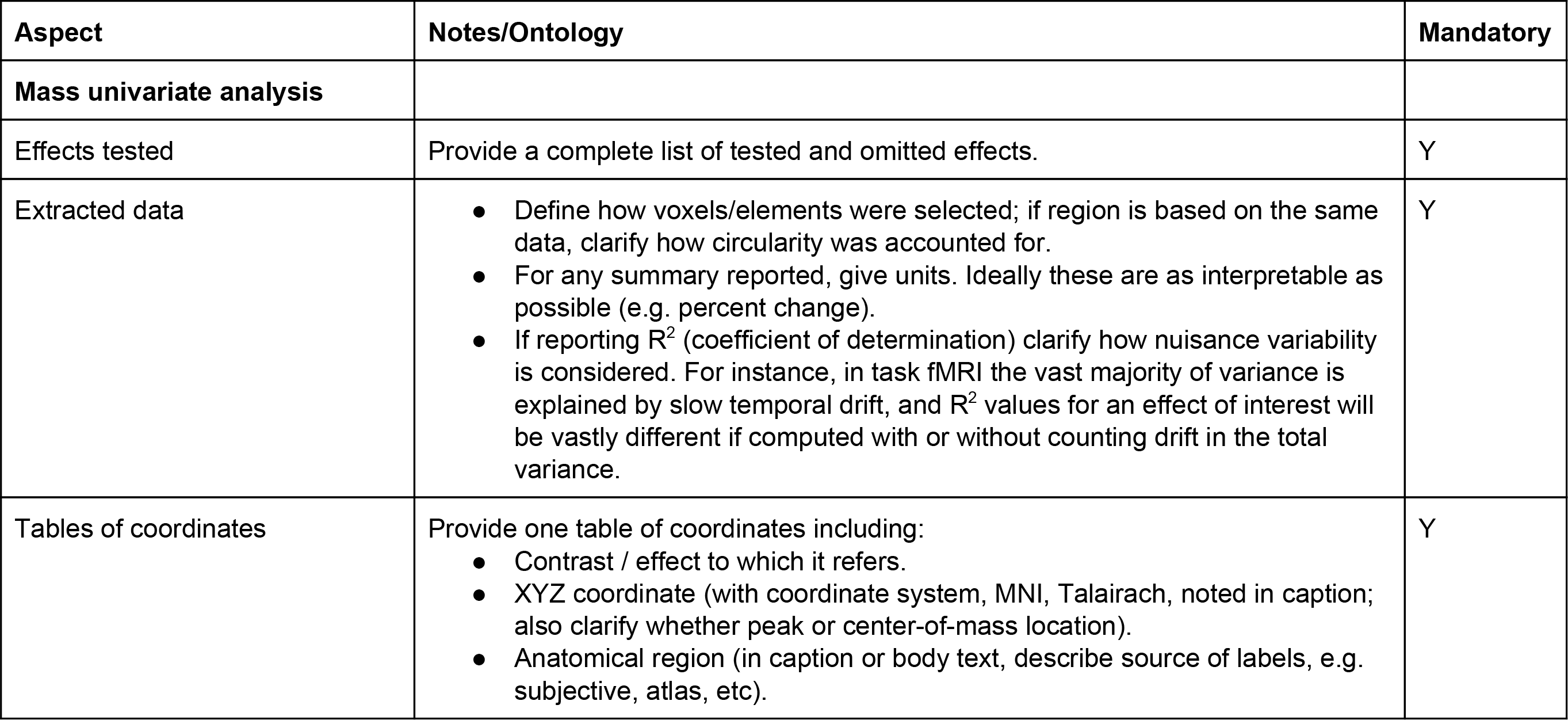
Results Reporting

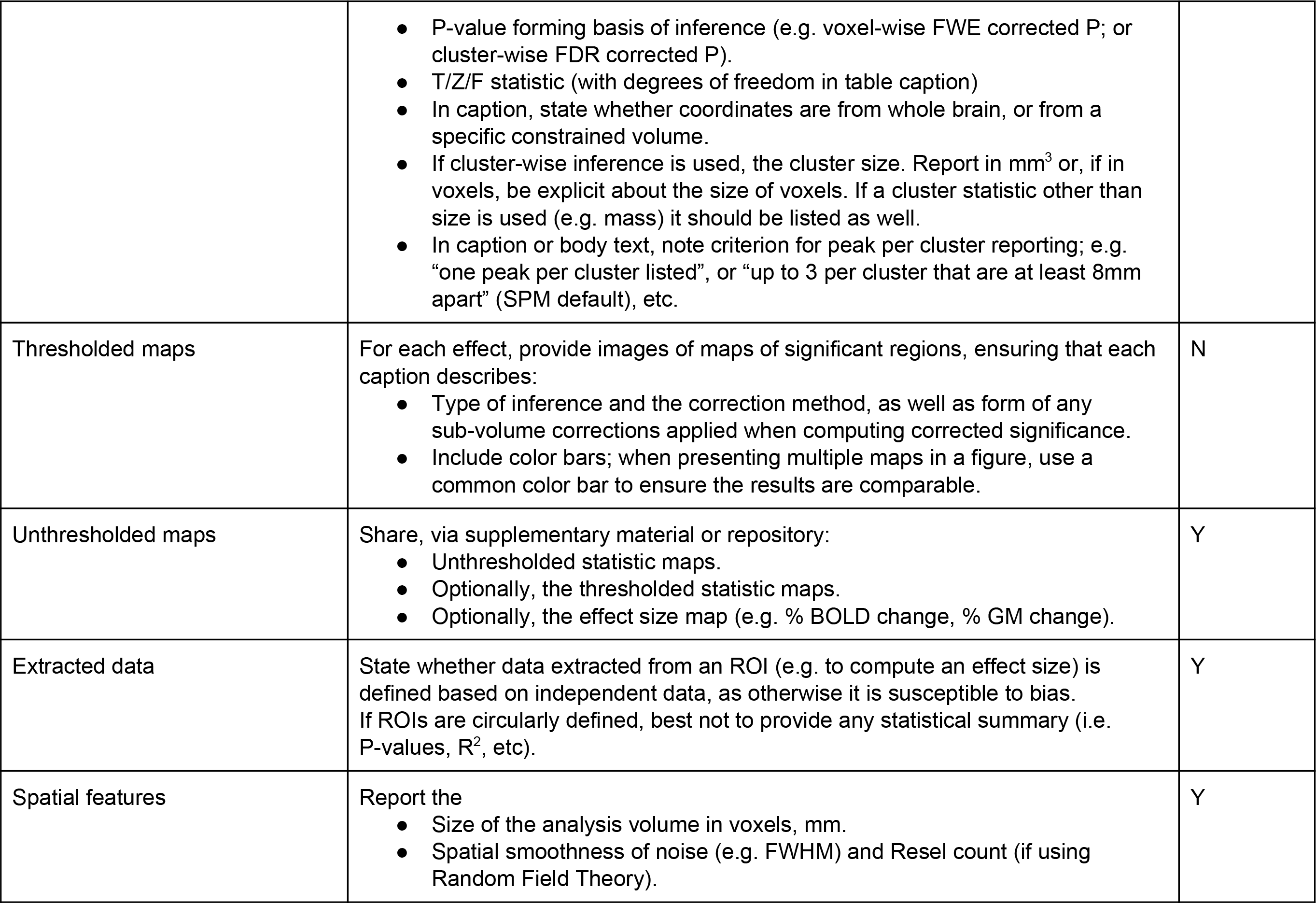

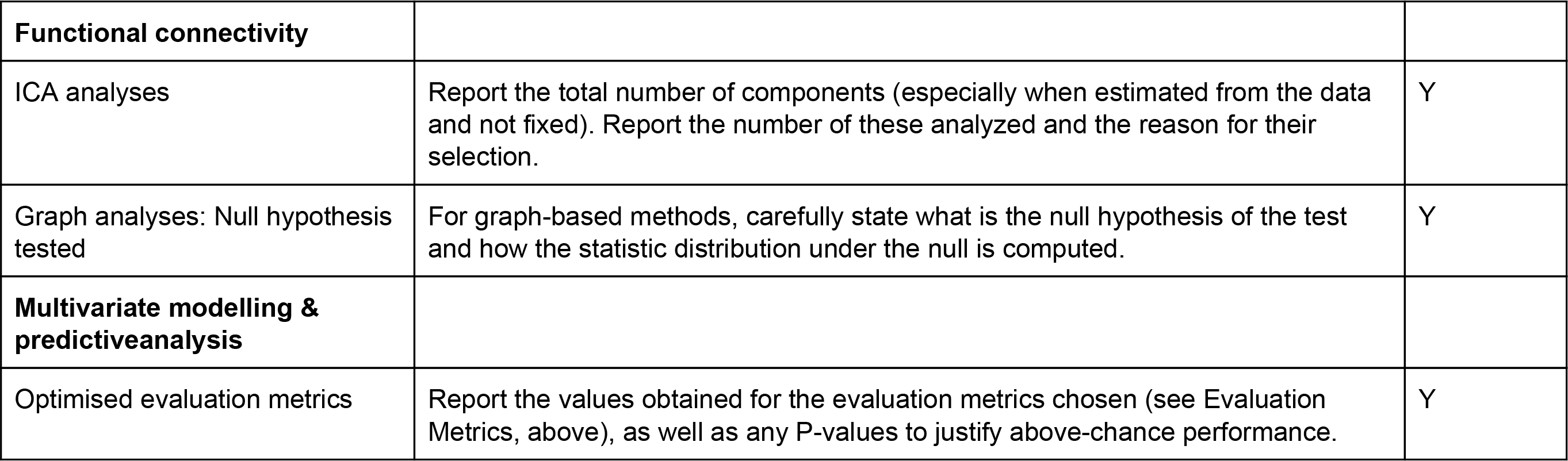

**Table D.6.**
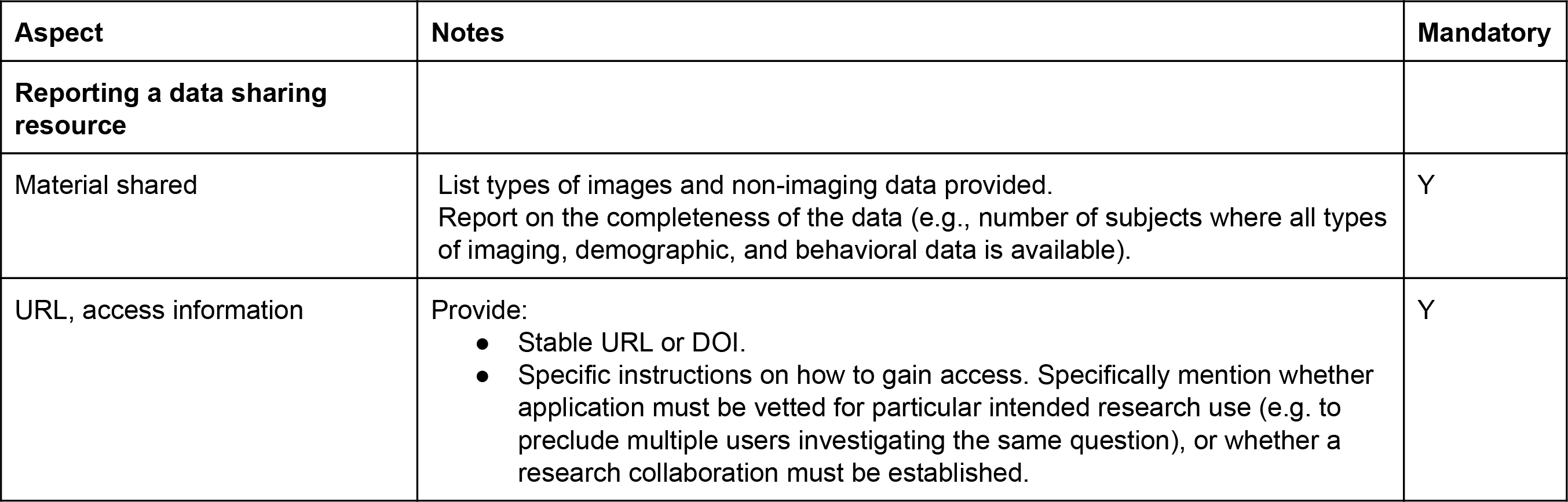
Data Sharing

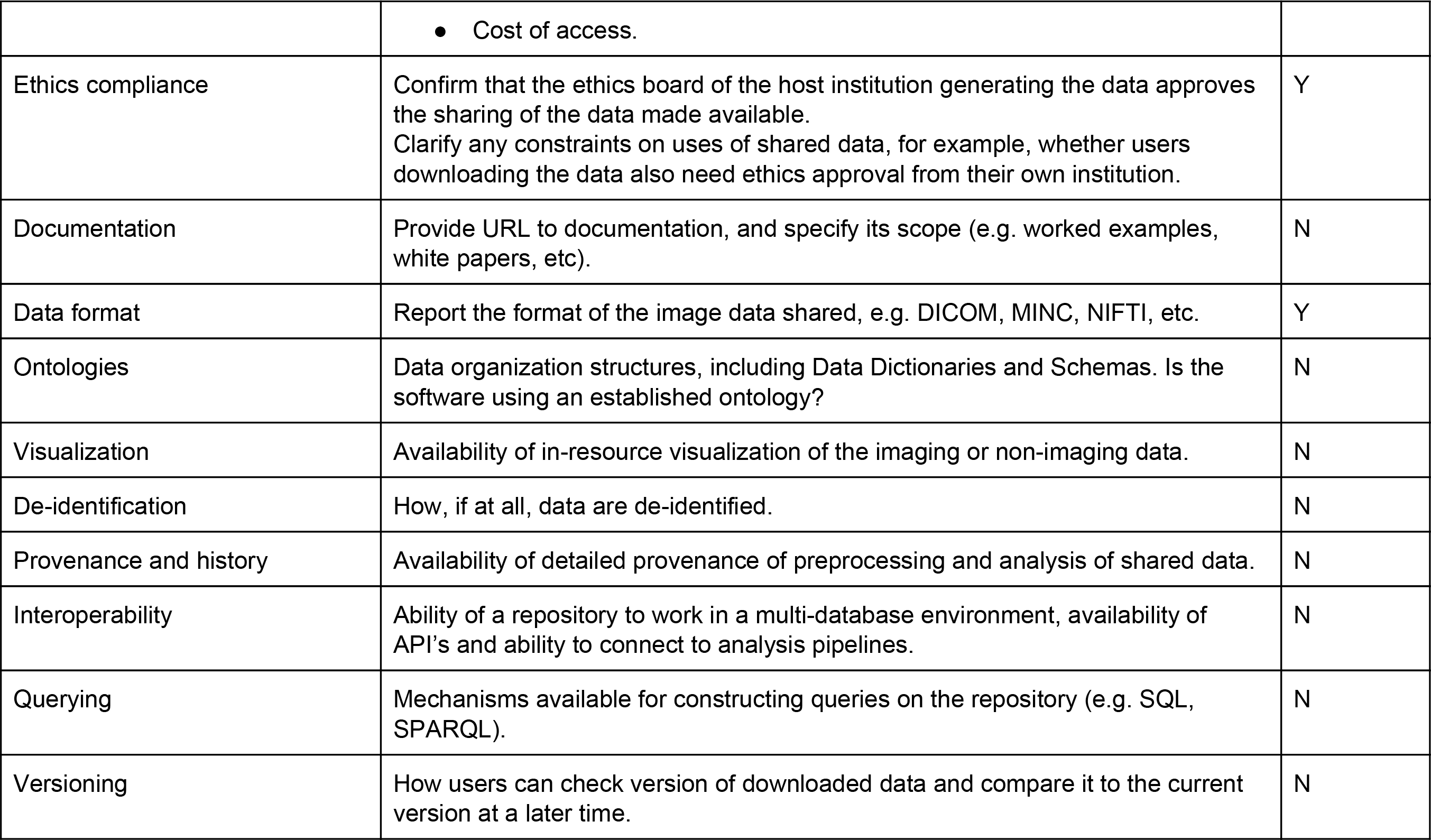

**Table D.7.** Reproducibility

| **Aspect** | **Notes/Ontology** | **Mandatory** |
| --- | --- | --- |
| **Documentation** |  |  |
| Tools used | Tool names, versions, and URLs. | Y |
| Infrastructure | Machine CPU model, operating system version, any use of parallelization. | Y |
| Workflow | Use of a workflow system, its version and URL. | N |
| Provenance trace | State whether detailed provenance information is available. | N |
| Literate program implementing results | Provide a URL linking to the relevant resource; for example, an ipython notebook implementing key analyses. | N |
| English language version | As the scientific lingua franca, documentation should be provided in English in addition to any other languages. | N |
| **Archiving** |  |  |
| Tools availability | Note if tools are publically available. | N |
| Virtual appliances | Note if a virtual environment to facilitate a repeated analysis is available. | N |
| **Citation** |  |  |
| Data | Provide permanent identifier if possible. | N |
| Workflow | Provide permanent identifier if possible. | N |

